# Systems genetics analysis identify calcium signalling defects as novel cause of congenital heart disease

**DOI:** 10.1101/2019.12.11.872424

**Authors:** Jose M.G. Izarzugaza, Sabrina G. Ellesøe, Canan Doganli, Natasja Spring Ehlers, Marlene D. Dalgaard, Enrique Audain, Gregor Dombrowsky, Alejandro Sifrim, Anna Wilsdon, Bernard Thienpont, Jeroen Breckpot, Marc Gewillig, Competence Network for Congenital Heart Defects, Germany, J. David Brook, Marc-Phillip Hitz, Lars A. Larsen, Søren Brunak

## Abstract

**Background:** Congenital heart disease (CHD) occurs in almost 1% of newborn children and is considered a multifactorial disorder. CHD may segregate in families due to significant contribution of genetic factors in the disease aetiology. The aim of the study was to identify pathophysiological mechanisms in families segregating CHD.

**Methods:** We used whole exome sequencing to identify rare genetic variants in ninety consenting participants from 32 Danish families with recurrent CHD. We applied a systems biology approach to identify developmental mechanisms influenced by accumulation of rare variants. We used an independent cohort of 714 CHD cases and 4922 controls for replication and performed functional investigations using zebrafish as *in vivo* model.

**Results:** We identified 1,785 genes, in which rare alleles were shared between affected individuals within a family. These genes were enriched for known cardiac developmental genes and 218 of the genes were mutated in more than one family. Our analysis revealed a functional cluster, enriched for proteins with a known participation in calcium signalling. Replication confirmed increased mutation burden of calcium-signalling genes in CHD patients. Functional investigation of zebrafish orthologues of *ITPR1*, *PLCB2* and *ADCY2* verified a role in cardiac development and suggests a combinatorial effect of inactivation of these genes.

**Conclusions:** The study identifies abnormal calcium signalling as a novel pathophysiological mechanism in human CHD and confirms the complex genetic architecture underlying CHD.

## Background

Congenital heart disease (CHD) represents malformations of the heart or intra-thoracic vessels, which affect cardiac function and occur in almost 1% of live births [1]. Most patients survive until adulthood and the number of adults with CHD has increased to above 3 million in Europe and the USA alone [2]. These patients are challenged by serious cardiovascular complications, which require specialized care, and their children have significantly increased CHD risk [3].

Although the specific cause of CHD is unknown for most patients, genetic factors contribute significantly to the aetiology (reviewed in [3, 4]).

Families presenting with recurrent cases suggest that mutations with large effects segregate with CHD in such families. Recent targeted next generation sequencing (NGS) efforts have shown that causative mutations may be identified in one third of CHD families [5]. In the majority of familial cases, the unidentified pathogenic variants may be explained by a "burden of genetic variation" model, which hypothesises that CHD may occur if the developing embryo is exposed to a certain burden of rare and common genetic variants, possibly in combination with epigenetic, environmental or stochastic effects [4].

Genomic information obtained by NGS analyses may further elucidate the genetic and molecular mechanisms causing CHD and thus contribute to improved health care and counselling of the patients and families. However, because of the complex genetic architecture of CHD, more sophisticated methods for interpretation of genetic variation needs to be developed before such information may be translated into clinical use.

Here, we performed whole exome sequencing (WES) in a cohort of 32 Danish families in which several family members presented with CHD. Utilising a systems biology approach, we discovered recurrent mutation of genes involved in calcium signalling. Loss-of-function analyses of three of the genes confirmed their crucial role in embryonic heart development.

## Methods

### Patient material

CHD families were identified through the Danish National Patient Registry (DNPR) and contacted by letter. Clinical diagnoses were validated by manual review of the patient files and detailed pedigrees were constructed based on interview with patients (Figure S1). DNA samples were obtained from a cohort consisting of 90 individuals in 32 families. Index patients were screened for mutations in the *NKX2-5* gene prior to WES [6]. Relatedness of family members was determined using the algorithm implemented in VCFtools. Pairwise relatedness of all 90 individuals is shown in Figure S2A. Relatedness between individuals not belonging to the same family peaks around zero, confirming that the families are unrelated. Individuals within families have positive relatedness score, confirming that they are related (Figure S2B).

The self-reported ethnicity of the individuals in our cohort was Danish. To verify the ethnic homogeneity of the 90 samples we used the algorithm for detection of outliers implemented in PLINK. For each individual in our cohort, we calculated the population distance to the 20 nearest neighbors using an identity-by-state (IBS) matrix based on 208,163 variants. These distances were compared to the mean of the population in terms of standard deviations (Z-scores). This test show that no samples have Z-scores below −2.5 confirming the ethnic homogeneity of the samples (Figure S3).

### Whole Exome Sequencing

Ninety exomes corresponding to 79 patients and 11 obligate carriers were sequenced. Capture followed the protocol corresponding to an Agilent Sure Select exome v4 kit and 100bp paired-end sequencing reads were generated on an Illumina Hiseq 2000 machine. Both exome capture and sequencing were performed at BGI Europe’s facility in Copenhagen. The average sequencing coverage was 91.8X. Eighty-six percent of the samples had a sequence coverage of 30X or higher in 80% of the exome. Cumulative sequencing coverage is shown in Figure S4A. Mean number of sequencing reads per sample was 64,396,134 (range 47,821,039-85,327,021). The distribution of sequencing reads is shown in Figure S4B.

Variants were stored in a mySQL database to facilitate filtering and comparison between both individuals and families. Population-wide allele frequencies were used to remove variants with a minor allele frequency higher than 1%. Finally, variants with unclear impact on gene function were removed (see definitions below).

### Bioinformatics pipeline for variant calling

Bioinformatics analysis of the sequencing data followed standard practices in the field and included assessment of data quality with FastQC; mapping to the human reference genome (hg19/Grhc37) with BWA mem [7]; removal of PCR duplicates, local indel realignment, base quality score recalibration, and variant calling with HaplotypeCaller, were performed with GATK [8]. Only variants with Phred scores ≥30 were considered. The functional consequences of variants were assessed with Ensembl’s Variant Effect Predictor (VEF) tool [9]. This step included the prediction of the pathogenicity of identified variants with both SIFT [10] and Polyphen-2 [11]. Variants were stored in a mySQL database to facilitate filtering and comparison between both individuals and families.

### Filtering of variants

Following genotype calling, variants were filtered to meet a number of sequencing quality requirements prior to consideration. The variants should correspond to single nucleotide events both in the reference and alternative alleles, be supported by read depth of at least 30 at the genomic position and have a Phred call quality greater than 30. Similarly, variants were required to be present in all affected members of a family, either in heterozygosity or in homozygosity.

Population-wide allele frequencies were used to further filter the candidate variants under the prerogative that the observed incidence of CHD is not coherent with highly frequent variants being causative factors. Variants were filtered from our analysis if they were observed with a minor allele frequency (MAF) higher than 1% in either any of the main populations (AFR, AMR, ASN, CEU) comprised by the 1000 Genomes project[12], the catalogue of Exome Annotation Consortium[13] (ExAC) or its European (excluding Finland) subpopulation. Similarly, we exploited the Danish ancestry of the patients for further reduction of the number of candidate variants. First, we discarded from further analyses those variants with MAFs higher than 1% in 2000 Danish exomes [14]. Second, variants present in 5% of the alleles of the 100 parents included in the Genome Denmark cohort [15, 16] were excluded in further analyses. This step implied a lift-over [17] of variants between the coordinates of reference genomes hg38 and hg19; in cases where multiple variants remapped to the same genomic position, the highest allele frequency for each possible allele was considered.

A final filtering exploited the functional consequences of variants according to affected genomic elements. Variants with unclear impact on gene function were disregarded. These included all noncoding variants (except intronic variants in splice-sites), stop codon variants, where the stop codon is retained and nonsense mediated decay (NMD) transcript variants.

### Enrichment of known CHD genes

We curated two lists of known CHD genes. A list of 829 genes which cause CHD when mutated in mice was derived from data in the Mouse Genome Informatics database (Table S1). A list of 144 human CHD disease genes was curated from the literature (Table S2). Statistical significance of overlaps was calculated using one tailed Fisher’s exact test, with a significance level of P<0.05.

### Definition of enriched protein-protein interaction modules

Protein-protein interactions (PPIs) were obtained from InWeb (InWeb5.5rc3), a scored high-confidence PPI database which contain 87% of human proteins and more than 500.000 interactions [18]. PPIs were represented as a network; nodes represent proteins and edges represent interactions between these proteins. InWeb contains protein interactions for 1,310 of the 1,785 candidate genes identified in our families. We used data from InWeb to generate a PPI network of these 1,310 genes and their first-degree interactors. The PPI network was pruned to include only high confidence relationships; we disregarded high-throughput yeast-2-hybrid experiments (“Matrix” interactions according to InWeb’s terminology) and interactions with a confidence below 0.1. After confidence filtering, the PPI network included a total of 8,186 genes and 29,463 interactions.

The PPI network was partitioned into overlapping modules (clusters) with ClusterOne [19] using the confidence score derived from InWeb to weight the clustering. Clusters were considered significant when below a corrected one-sided p-value of 0.05 and a minimum of five proteins. Highly connected clusters of proteins identified in this fashion constitute a proxy for functionally related protein complexes.

Enrichment of candidate genes was determined for each cluster using a permutation analysis (k=10e4). Correction for multiple testing was performed using the procedure of Bonferroni-Holm. A one-sided p-value of 0.05 was used as significance level to identify clusters significantly enriched for candidate genes.

Probability of being loss-of-function intolerant (pLI) values of genes encoding the 27 proteins in the calcium signalling module, 829 and 144 known CHD genes from mouse models and patients, respectively, were obtained from the ExAC database. The distributions were compared to all 18,225 genes listed in the ExAC database using a Kruskal-Wallis One Way Analysis of Variance on Ranks. Significance level was 0.05.

### Replication

WES data from a previously published, independent cohort [20] was used for replication; WES data from a total of 714 non-syndromic CHD cases and 4922 controls of European ancestry was analyzed.

Control samples were participants in the INTERVAL randomized controlled trial were recruited with the active collaboration of NHS Blood and Transplant England, which has supported field work and other elements of the trial. DNA extraction and genotyping was funded by the National Institute of Health Research (NIHR), the NIHR BioResource and the NIHR Cambridge Biomedical Research Centre. The academic coordinating center for INTERVAL was supported by core funding from the NIHR Blood and Transplant Research Unit in Donor Health and Genomics, UK Medical Research Council (G0800270), British Heart Foundation (SP/09/002), and NIHR Research Cambridge Biomedical Research Centre. Replication was performed by comparing the number of cases and controls harboring very rare (MaF<0.001) variants in any of the genes *ADCY2*, *ADCY5*, *CACNA1D*, *CACNA1H*, *CACNA1I*, *CACNA1S*, *GRIA4*, *ITPR1*, *NFAT5* and *PLCB2*. Data from *CACNA1F* was missing, thus this gene was excluded from the analysis. Synonymous variants, protein altering variants, PAV (inframe_insertion, start_lost, stop_retained, stop_lost, missense, inframe_deletion, protein_altering, start_retained) and protein truncating variants, PTV (stop_gained, splice_donor, splice_acceptor, frameshift) were identified using the Variant Effect Predictor tool (VEP API v90). Quality control and filtering were performed using Hail 0.2. In brief, both samples and variants were included for further analysis if they met the following filtering criteria: call rate >= 0.95, genotype quality average >= 25 and depth average >= 15. In addition, only Genotypes (heterozygous) with allelic balance (AB) within the range 0.25-0.75 were retained in the analysis. Deleterious variants were identified by assigning a Combined Annotation Dependent Depletion (CADD)-score [21] and a score based on regional missense constraint (MPC-score) [22]. MPC score >2 was used to identify pathogenic variants. Previous analysis of *de novo* variants identified in 5620 cases with neurodevelopmental disorders shoved that variants with MPC>2 have a rate 5.79 times higher in cases than controls. Statistical significance was calculated using two-tailed Fisher’s exact test, with a significance level of P<0.05.

To test the specificity of the observed burden of rare variants we created 10,000 random sets of 10 genes with the same size distribution as the gene-set of 10 calcium signalling genes, which were analyzed in the replication study. For each of the 10,000 random gene-sets we counted the number of cases with pathogenic variants (MPC score >2).

### Zebrafish maintenance and microinjection

AB/TL zebrafish strain (obtained from the Zebrafish International Resource Center) was used in the experiments. Embryos were maintained and staged as previously described [23, 24]. 1.5, 3, 6 ng of adcy2a-SP-MO (5’-GGATGAGGGTAACTCACCTGACATT-3’), itpr1b-SP-MO (5’-GTGCATAAACGCGGCCTTACCTCGA-3’), plcb2-SP-MO (5’-CTGTAGTTTCTGTTCACCTCATCAG-3’), standard-MO (5’-CCTCTTACCTCAGTTACAATTTATA-3’) and 0.5, 1, 2 ng of adcy2a-SP-MO2 (5’-CCCCCAGTCTCCAAACACTCACCAG-3’), itpr1b-SP-MO2 (5’-CCAGACTGTAGACAAGAGAGACATG-3’) and plcb2-SP-MO2 (5’-TGTGGTAAAGGATACTCCACCCAGT-3’) (Gene Tools, LLC) were injected into one-cell stage embryos and harvested at 48 hpf. Embryos were imaged under Zeiss AxioZoom V16 (Carl Zeiss, Brock Michelsen A/S, Denmark).

In order to verify SP-MO efficiencies embryos injected with SP-MOs were collected at 48 hpf and RNA was isolated by QIAzol reagent (Qiagen). cDNA synthesis was performed using iScript Reverse Transcription Supermix (Bio-Rad). SP-MO knock-down were assessed by PCR using gene specific primers (Table S3) spanning the SP-MO-targeted exon.

### Whole-mount In Situ Hybridization

Digoxigenin (DIG)-labelled anti-sense *myl7* and *mef2cb* [25] riboprobes were synthesized from linearized pGEM-T easy and pCMV-SPORT6.1 vectors respectively using the DIG RNA labeling mix (Roche) and the T7 RNA polymerase (Roche). Embryos collected at 48 hpf were raised in the presence of 0.2 mM 1-phenyl-2-thiourea (PTU) upon gastrulation for optical clarity [24]. For analysis of *myl7* and *mef2cb* expression, embryos at 48 hpf and 10 somite stage, respectively, were dechorionated and fixed in freshly prepared 4% paraformaldehyde in phosphate buffered saline (PBS, pH 7.4) overnight. Whole-mount in situ hybridization was performed as previously described with minor modifications [26]. Embryos were imaged under Zeiss AxioZoom V16 (Carl Zeiss, Brock Michelsen A/S, Denmark). The staining intensity of *mef2cb* riboprobe was measured as integrated density (IntDen, ImageJ software-NIH, USA) and mean values from three embryos of each group were plotted relative to wild type value. Data show mean±std dev.

### Real-time quantitative RT-PCR

Total RNA from pools of 50 zebrafish embryos was extracted using Trizol (Ambion) and a RNeasy mini kit (Qiagen) and used to synthesize random-primed cDNA (SuperScript II Reverse Transcriptase, Invitrogen). A Brilliant III Ultra-Fast SYBR® Green QPCR Master Mix (Agilent Technologies) was used for cDNA amplification. Samples were analyzed using a 7500 fast real-time PCR system (Applied Biosystems). Data were normalized to the average expression of housekeeping genes *actb1* and *rpl13a*. RT-PCR primers are listed in Table S3.

## Results

### Exome sequencing data analysis

We performed whole exome sequencing of 90 individuals belonging to 32 multiplex CHD families (Figure S1-S4). To identify potentially disease-causing genes, we analysed rare variants shared by affected family members. The analysis was performed after removing the variants that are likely sequencing artefacts, that occupy genomic elements with a mild functional consequence or that can be found widespread in the general population (materials and Methods section, workflow shown in Figure S5). We identified 3,698 rare inherited variants in 1,785 genes, denoted candidate disease genes (CDGs) hereafter. The number of CDGs per family ranged from 56 to 507 (Figure S6).

### Recurrent candidate disease genes

To test if particular candidate genes were overrepresented, we calculated the fraction of CDGs shared by all possible pairs of families in the cohort (Figure 1A, Figure S7). Pairs of families only share a small fraction of their CDGs (Median = 0.05, 72.3% of pairwise values <0.2). The low overlap of CDGs between pairs of families and the absence of major disease genes across families suggest that each family has a unique constellation of disease genes, in line with the complex polygenic aetiology of CHD.

**Figure 1.**
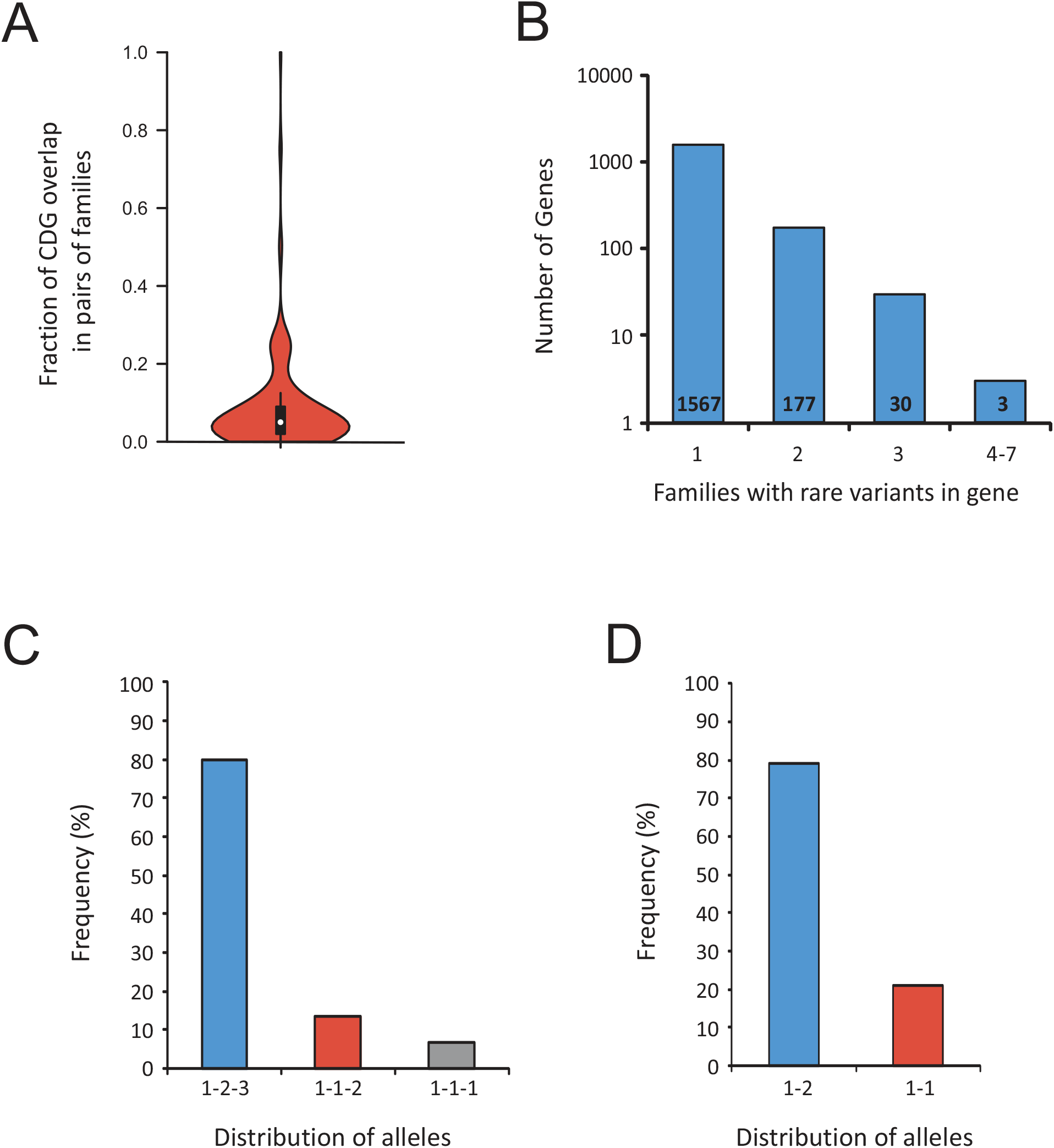
Distribution of candidate disease genes and variants across families. (A) Overlap between CDGs in pairs of families. (B) Distribution of CDGs across families. The number of CDGs found in one, two, three and 4-7 families is shown. (C) Distribution of alleles in CDGs found in three families (1-2-3: three different alleles, 1-1-2: two different alleles, 1-1-1: same allele found in all three families). (D) Distribution of alleles in CDGs shared in two families (1-2: different alleles, 1-1: same allele).

To analyse this in more detail, we calculated the number of families with rare inherited variants in each of the 1,785 CDGs. None of the 1,785 CDGs was mutated in more than seven families (Figure 1B, Figure S8) after disregarding genes that rely on the accumulation of variants to exert their biological function, like hypermutated genes in the MHC machinery or genes encoding olfactory receptors.

We identified 218 CDGs shared between two or more families. For variants shared between two and three families, less than 20% were identical across families (Figure 1 C-D). We propose that these variants might partially explain the origin of the disease. For example, three different rare alleles of *DNAH5* were identified in affected members of families 489, 732 and 1121, where patients presented with ASD, VSD and outflow tract defects. Mutation of *DNAH5* is associated with Primary Ciliary Dyskinesia (PCD, OMIM #608644) [27]. A subset of PCD patients present with CHD [28]. Another example of a CDG where rare variants were found in more than one family is *KMT2D*, encodes a histone methyltranferase involved in heart development [29]. Mutation of *KMT2D* is associated with Kabuki syndrome (OMIM #147920), a rare developmental disorder which includes CHD as part of a wide phenotypical spectrum [30].

### CDGs are enriched for known disease genes

We expect that a subgroup of variants in our 1,785 CDGs could be causative mutations leading to CHD. In such a scenario, a diverse but limited number of genomic variants, acting by the accumulation of additive effects, could explain the occurrence of CHD in individual families. This etiological diversity at the genomic level would hinder detection with traditional association analysis, but we hypothesized that the CDGs would be enriched for known CHD disease genes. To test this, we calculated the overlap between the 1,785 CDGs in our families and curated lists of genes known to cause CHD in mouse models and patients, when mutated (Table S1 and S2). We observed significant enrichment of known CHD genes among CDGs affecting five or fewer families (Figure 2A and B). When only variants scored pathogenic by SIFT/Polyphen-2 are considered, this enrichment increases Figure S9). In order to validate the CDG list and explore the possibility of a selection bias in our sets of known CHD genes, we produced 10,000 random gene sets (one for each of the two sets) and ascertained the overlap with the CDGs. This analysis corroborated the statistical significance of the observations (p-value<0.0001 for both sets).

**Figure 2.**
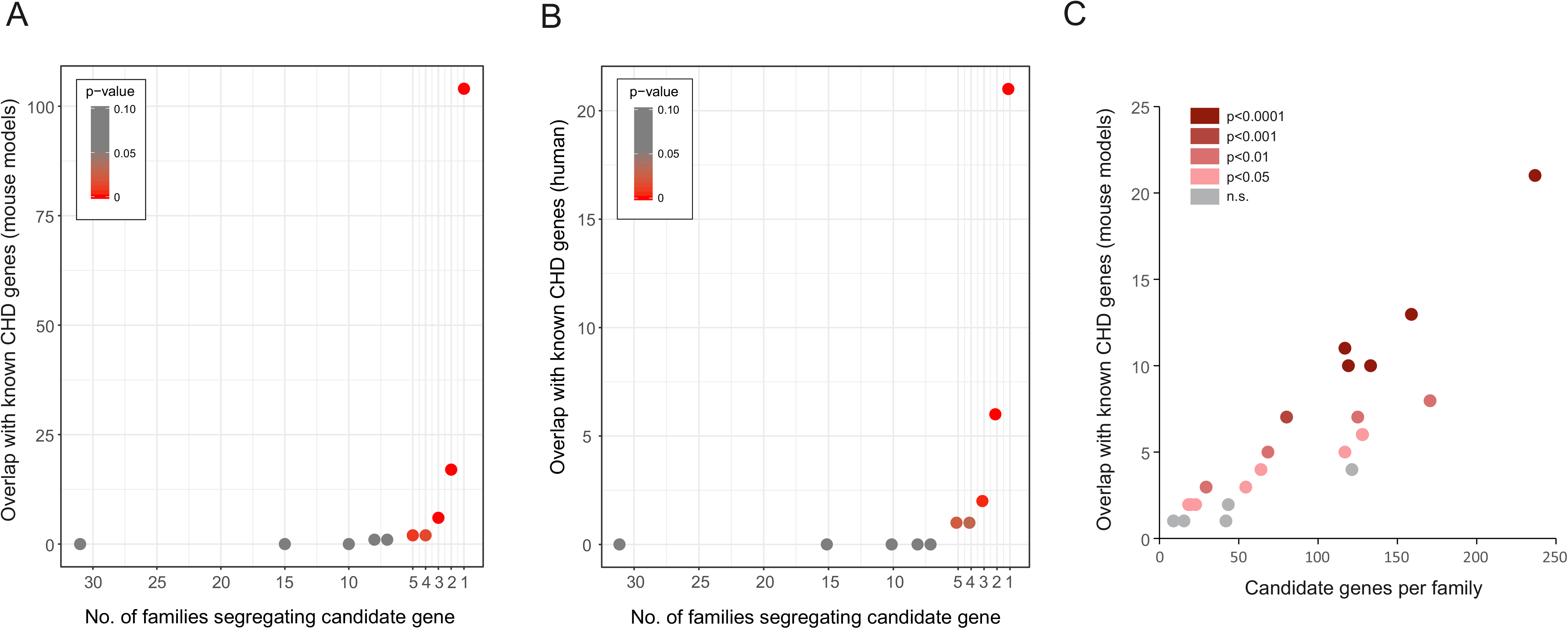
Enrichment of CHD genes in candidate disease genes. Overlap between CDGs and known CHD genes from mouse models (A) and patients (B). The number of overlapping genes is plotted against the number of families the CDGs were found in. (C) The number of overlapping genes (mouse models) per family. Statistical significance of the overlap is indicated by color code (red colors: significant, grey color not significant, n.s.).

Distribution of known genes across families is shown in Figure S10 and Table S4. Individual families often present with rare mutations in more than one known CHD gene, suggesting a substantial fraction of the observed heart defects might be explained by a combination of rare mutations inherited within the families. Under the hypothesis that CHD runs within individual families as the result of such a combined effect from mutations in several developmental genes, we would expect that families present significant enrichment of mutations harboured by known CHD genes. Enrichment of CHD genes from mouse models per family was determined using a permutation test (n=10,000). Affected individuals from 23 families share rare mutations in known CHD genes. In 19 of these 23 families (78.3%) enrichment of known CHD genes among the CDGs is statistically significant at p<0.05 (Figure 2C).

### Rare variants affect functional modules in a protein-protein interaction network

We further investigated whether CHD was caused by the disruption of cooperative protein functionality at the systems level rather than at the individual gene level. A protein-protein interaction network was generated using 1,310 seed genes (the subset of the 1,785 CDGs for which InWeb has recorded protein interactions). Their first-degree neighbours and their interactions from the current version of InWeb (InWeb5.5rc3) [18] were added as well, summing up to a total of 8,186 proteins and 29,463 interactions. Using the graph clustering algorithm ClusterOne [19], we identified 230 significant PPI clusters considering a threshold p-value of 0.05 and a minimum number of five genes. Of these, 25 clusters presented two or more genes mutated in at least one of the families (Table S5). By permutation analysis (k=10,000) we identified two clusters accommodating more CDGs than expected by chance (p-values of 0.039 and 0.0033, respectively).

One of these corresponds to a small module, encoded by of five genes (*INSL1*, *RXFP2*, *RLN1*, *RLN2*, *RXFP1*) where the three first are CDGs. Due to the small size and low connectivity of this cluster, we decided to focus on the second cluster.

The second significant cluster corresponds to a highly interconnected group of 27 proteins, encoded by eleven CDGs and their first-degree interaction partners (Figure 3A). These CDGs present mutations in ten different families. Affected individuals from three families shared rare mutations in more than one of the eleven genes, and two genes (*ITPR1* and *CACNA1S*) were each mutated in two different families (Table S6).

**Figure 3.**
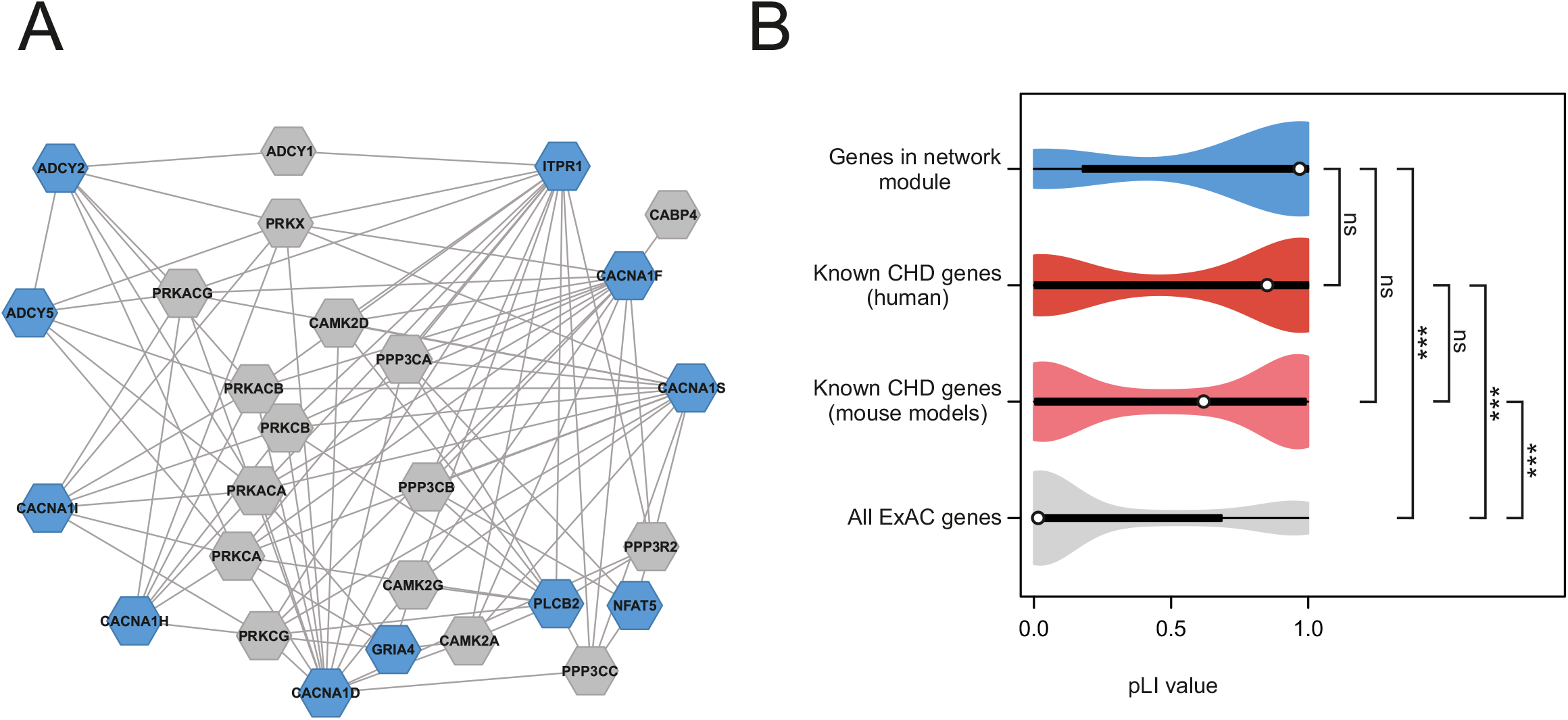
Identification of a calcium signalling network affected by rare mutations identified in CHD families. (A) Network module of CDGs (blue) and their interaction partners (grey). The module accommodates more CDGs than expected by chance (adjusted p-value 0.0033). Proteins are shown as hexagons, protein interactions are shown with lines. (B) Violin plots of distributions of pLI scores in genes encoding the 27 proteins in the network module (upper, blue), known CHD genes from patients and mouse models (middle, red and pink, respectively) and all 18,225 genes listed in ExAC with a calculated pLI score (lower, grey). ***:P<0.001. ns=not significant (Kruskal-Wallis One Way Analysis of Variance on Ranks).

The 27 proteins in the cluster constitute calcium channels or signal transduction enzymes such as adenylate cyclase that interacts with calcium dependent protein kinases. A Gene Ontology term enrichment analysis using AmiGO [31] for the 27 proteins in the cluster showed enrichment in biological processes involved in calcium signalling (Table S7).

To investigate the functional importance of the genes encoding proteins in the cluster, we compared the probability of being loss-of-function intolerant (pLI) of these 27 genes with known CHD genes and all 18,225 genes listed in the Exome Aggregation Consortium database. The distribution of pLI scores of known CHD genes have median pLI values of 0.63 and 0.86, respectively, and the distributions suggest that more than half of known CHD genes are intolerant against loss-of-function mutations (Figure 3B). The distribution of pLI scores of the 27 genes encoding proteins in the significant cluster is skewed towards the higher values of the distribution (median = 0.98), suggesting intolerance to loss-of-function mutations and supporting the hypothesis that these genes might potentially play an active role in the aetiology of CHD.

### Replication

Using WES data from an independent cohort of 714 CHD cases and 4922 controls [20], we tested the mutation burden of the same calcium-signalling gene-set, as we found mutated in our families (genes listed in Table S6). In this gene-set, we analysed the distribution of CADD and MPC variant scores between CHD cases and controls, and observed significant different score-distributions of rare (MAF < 0.001) variants, predicted to alter the gene products (i.e. PAV and PTV)(Figure 4). Variants with MPC score above 2 (MPC2 variants) have been shown to be significantly associated with pathogenesis [22]. Thus, to test for burden of pathogenic mutations, we calculated the number of cases and controls harboring MPC2 variants in the calcium signalling gene-set. We observed more than 2-fold enrichment (OR 2.68, P-value 3.7e-04) of such variants in CHD cases (Table S8). To test if this enrichment was specific for the calcium-signalling gene-set, we created 10,000 random gene-set with the same size-distribution as the calcium genes. For each random set, we counted the number of CHD cases harboring MPC2 variants. Only four of the random gene-sets were found with more MPC2 variants in cases than the calcium-gene-set (Figure S11). These results confirm an increased burden of pathogenic mutations in genes involved in calcium-signalling among CHD patients.

**Figure 4.**
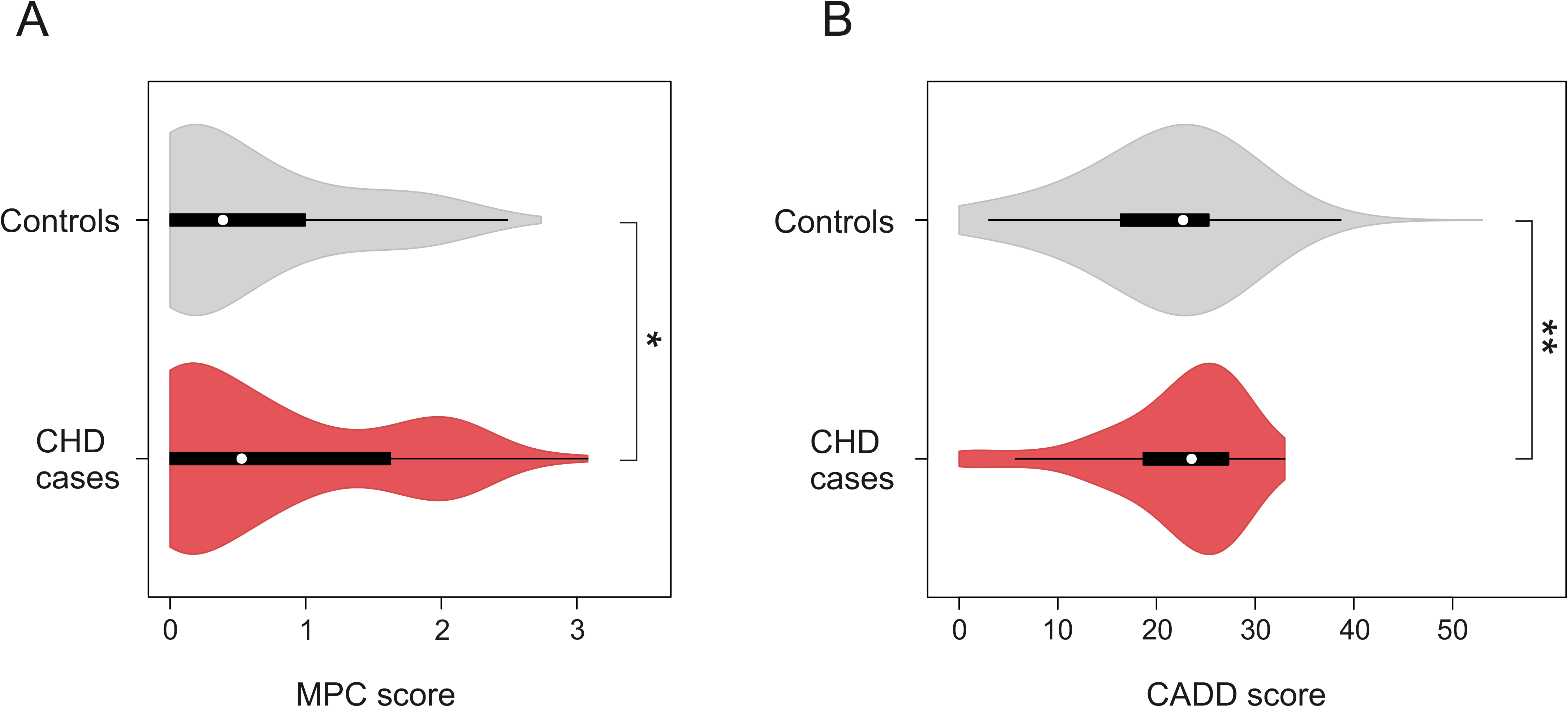
Distribution of MPC and CADD Scores of rare variants in 714 CHD Cases and 4922 Controls. Protein altering and truncating variants (PAV and PTV) with MAF < 0.001 identified in the genes *ADCY2*, *ADCY5*, *CACNA1D*, *CACNA1H*, *CACNA1I*, *CACNA1S*, *GRIA4*, *ITPR1*, *NFAT5* and *PLCB2* were scored using MPC score [22] (A) or CADD score [21] (B). N_CHD_ =136 variants. N_Controls_ = 982 variants. Difference between median values of controls and cases was determined using a Mann-Whitney Rank Sum Test. **: P<0.01, *: P<0.05.

### Knock-down of *adcy2a*, *itpr1b* and *plcb2* cause cardiac malformations in zebrafish

We used zebrafish as a model to address the functional relevance in heart development, of genes within the identified cluster. In the cluster, 11 out of 27 genes were mutated in the families we assessed. From these 11 genes, we assessed the consequence of loss-of-function for three zebrafish genes, *adcy2a*, *itpr1b*, *plcb2*. These genes are orthologues to the human genes *ADCY2*, *ITPR1* and *PLCB2*, encoding calcium signalling proteins, of which roles in cardiac development have been elusive.

We found that knock-down of either of the three genes by morpholino oligonucleotides gave rise to abnormal morphology or laterality of the heart (Figure 5A). The abnormal morphology included aberrant atrioventricular canal (AVC) formation where ring structure of AVC was hindered, mis-looping of the heart and narrowing in atrium or ventricle. Laterality defects reflected straight or reversed hearts. Injection of *adcy2a*-MO, *itpr1b*-MO and *plcb2*-MO resulted in cardiac defects in 53%, 73% and 66% of embryos, respectively (Figure 4B).

**Figure 5.**
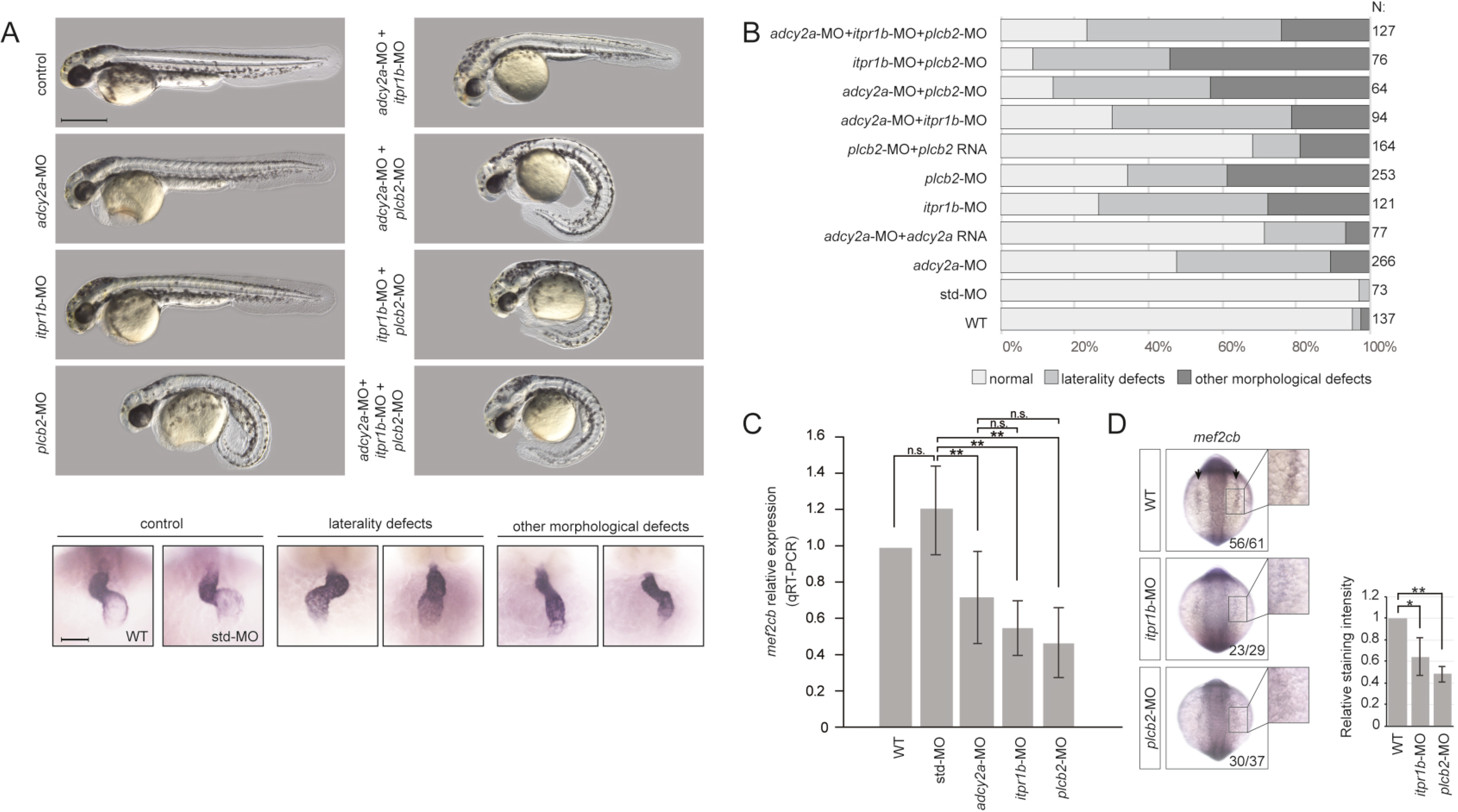
Functional validation of candidate genes in zebrafish. (A) Phenotype of controls and morphants. Single genes and combinations of genes targeted by splicing morpholinos are indicated on the left. Upper panels: gross appearance of 48 hpf zebrafish embryos. Lower panel: examples of cardiac phenotypes of 48 hpf wt, control and morphant embryos. Hearts were visualised by ISH with a probe against the cardiac marker *myl7*. (B) Quantification of phenotypes in wt, controls, morphants and mRNA rescued morphants. Note the combinatorial effects on cardiac phenotypes when more than one gene is affected. (C, D) Expression of *mef2cb* in 10 somite stage zebrafish embryos, analyzed by qRT-PCR (C) and ISH (D). ISH staining intensity was quantified and analyzed using student’s T-test. *: P<0.05, **: P<0.01.

In addition to the cardiac defects, knock-down of *itpr1b* causes edema in the brain, whereas curved tail was seen by knock down of *plcb2* (Figure 5A).

Knock-down of more than one gene by injections of both efficient and sub-efficient doses of MO resulted in an increase of the number of embryos with heart defects, suggesting a combinatorial effect (Figure 5B, Figure S12).

We showed that the morpholinos work efficiently causing splicing defects (Figure S13) and we validated the specificity of the morpholinos, by using a second set of morpholinos and co-injection of *in vitro* expressed mRNAs encoding wildtype proteins (Figure 5B, Figure S14). Unfortunately, we were not able to clone the cDNA of *itpr1b*, presumably due to the large size of the transcript (8.4 kbp coding region), but co-injection of wildtype *adcy2a* and *plcb2* mRNA resulted in partial rescue of the cardiac phenotype, confirming the specificity of the morpholinos.

To investigate effects of gene knock-down at the molecular level we analysed the expression of *mef2c*, a transcription factor which is involved in cardiac morphogenesis and myogenesis and known to be a specific target for Ca^2+^ dependent control of gene expression in cardiomyocytes [32, 33]. In zebrafish two copies of *mef2c* exists. We analysed the expression of *mef2ca* and *mef2cb* in wild type, control and morphant embryos at 10 somite stages, where both genes are expressed in the bilateral heart fields at the anterior lateral plate mesoderm. We observed significantly reduced expression of *mef2cb* in *adcy2a*, *itpr1b* and *plcb2* morphants (Figure 5C and D).

## Discussion

Interpretation of variants detected in exomes from CHD patients is challenging due to the extreme heterogeneity which characterise CHD [4]. Analysis of complex networks has previously been applied successfully to interpret genetic variants identified in CHD patients [34–36]. Here, we used a scored human PPI network [18] to interpret rare variants identified in Danish families with CHD.

We analysed the exomes of 90 individuals in 32 multiplex CHD families for rare variants which were shared among affected individuals within a family and identified a total of 1,785 potential CHD disease genes. The large number of candidate genes, underlines the challenge in interpreting variants in CHD patients, even in multiplex families. However, our analysis showed that the 1,785 CDGs were enriched for genes known to cause heart defects.

Approximately 12% of the CDGs were found in more than one family, but none of them was represented in more than seven families, and on average less than 10% of the CDGs found in a family were shared with another family. Thus, our data support that CHD is an extremely heterogeneous disorder and consistent with a model, where several rare genetic variants contribute to the pathogenesis in the individual patient.

We integrated PPI data in our analysis of CDGs and discovered that eleven of the CDGs converge in a functional cluster of genes, which encode proteins involved in calcium signalling. Analysis an independent case-control cohort confirmed increased mutation burden of this calcium-signalling gene-set in CHD patients.

Intracellular calcium plays essential roles in physiology and pathophysiology of the adult heart. In the healthy heart, intracellular calcium fluxes control cardiomyocyte contraction and mutations in calcium handling genes may cause arrhythmia [37]. In addition, calcium also modulate transcription in cardiomyocytes through a complex signalling network, and abnormal calcium handling and signalling is part of the pathophysiology in congestive heart failure [32, 38].

Defects in calcium signalling have not previously been associated with CHD in humans, but animal studies implicate an important role of calcium signalling in heart development. Pharmacological blockade of L-type calcium channels during embryonic development cause heart defects in mice ^28^. Targeted deletion of *Nfatc1* is embryonic lethal in mice and causes malformation of the cardiac valves, outflow tract and ventricular septum [39, 40]. Overexpression of activated calcineurin rescues cardiac developmental defects in calreticulin deficient mice [41].

Two of the 27 proteins within the cluster we identified were previously implicated in cardiac development and CHD. Knockout of *Nfat5* is embryonic lethal in mice and results in reduced compaction of cardiomyocytes in the ventricular wall and trabeculae [42]. Double knockout of *Itpr1* and *Itpr3* results in aberrant development of the outflow tract and right ventricle, while mice defective of only one of the two genes develop normally [43]. Likewise, double knockout of *Itpr1* and *Itpr2*, is embryonic lethal and results in cardiac malformation [44], together suggesting a redundant role of IP_3_ receptors in mammalian heart development.

Functional validation of *ADCY2*, *ITPR1* and *PLCB2* in a zebrafish model confirmed a critical role during embryonic heart development. We observed defects in cardiac morphology and laterality in a significant proportion of embryos injected with morpholinos targeting *adcy2a*, *itpr1b* and *plcb2*. Importantly, we observed more embryos with cardiac defects when combinations of two and all three genes were knocked down, supporting that the three genes interact functionally in heart development. Specificity of the morpholinos used in the experiments was confirmed by rescue experiments and further corroborated by significant reduction in the expression of *mef2cb*, a zebrafish orthologue of *Mef2c*, which is a well-established target of calcium signalling in cardiomyocytes [43, 45].

The cardiac malformations, present in our family cohort and replication cohort, were unselected with respect to severity or anatomical similarity. Thus, our data indicate that calcium signalling defects are associated with both familial and sporadic CHD and unrelated to specific groups of malformations.

We have recently shown that specific malformations co-occur in families, suggesting that specific developmental programs may be responsible for certain groups of heart malformations [46]. We suggest that WES or whole genome sequencing of cohorts of families selected for specific malformations, combined with systems-based analysis, as presented here, may be a useful strategy for identification of pathophysiological mechanisms in CHD.

## Conclusions

Our data support a model where CHD is caused by a combinatorial effect of rare and common genetic variants. Our systems level analysis of rare genetic variants, segregating with CHD in families, and functional analysis in zebrafish, identified defects in calcium signalling as a novel pathophysiological mechanism in CHD.

## Acknowledgements

The mef2cb probe plasmid was a kind gift of Dr. Yaniv Hinits, King’s College London. We thank Matthew E. Hurles, Wellcome Trust Sanger Institute for providing access to published WES control data. Control samples were participants in the INTERVAL randomised controlled trial.

## Availability of data and materials

Exome sequencing data from Danish families are available upon request. Other data generated or analysed during this study are included in the main paper or its additional files.

## Authors’ contributions

SB and LAL designed and supervised the project. SGE established the family study cohort. MPH, JB, MG, DB and Competence Network for Congenital Heart Defects, Germany established the secondary patient cohort. JMGI, SGE, LAL, EA, MPH, AS, AW, BT, NSE, MD and GD generated and/or analysed the data. CD performed zebrafish experiments. All authors contributed to writing the manuscript and approved the final manuscript.

## Ethics approval and consent to participate

This project was approved by the Danish Data Protection Agency (2009-41-3570) and the Danish National Board of Health (H-D-2009-070). Health records were accessed only after explicit consent by the patients. All zebrafish research was approved by and conducted under license from the Danish Animal Experiments Inspectorate.

