## Supplemental data for "Systems genetics analysis identify calcium signalling defects as novel cause of congenital heart disease"

**Members of the Competence Network for Congenital Heart Defects, Germany**

Hashim Abdul-Khaliq, Hans-Heiner Kramer, Felix Berger, Brigitte Stiller, Ulrike Bauer, Thomas Pickardt, Sabine Klaassen

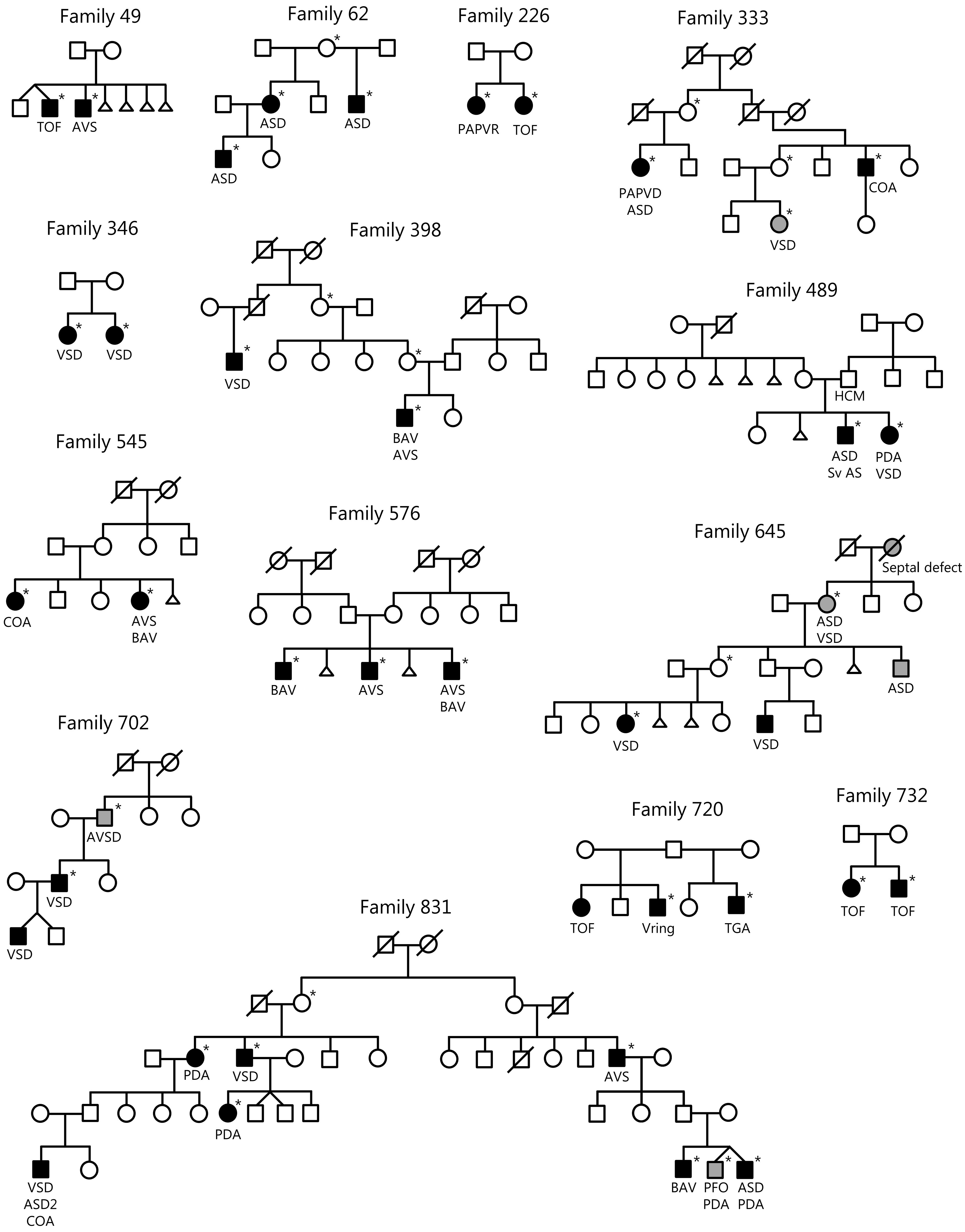
Family 49
TOF
AVS
Family 62
ASD
ASD
ASD
Family 226
PAPVR
TOF
Family 333
PAPVD
ASD
COA
VSD
Family 346
VSD
VSD
Family 398
VSD
BAV
AVS
Family 489
HCM
ASD
Sv AS
PDA
VSD
Family 545
COA
AVS
BAV
Family 576
BAV
AVS
AVS
BAV
Family 645
Septal defect
ASD
VSD
ASD
VSD
VSD
Family 702
AVSD
VSD
VSD
Family 720
TOF
Vring
TGA
Family 732
TOF
TOF
Family 831
PDA
VSD
AVS
PDA
VSD
ASD2
COA
BAV
PFO
PDA
ASD
PDA

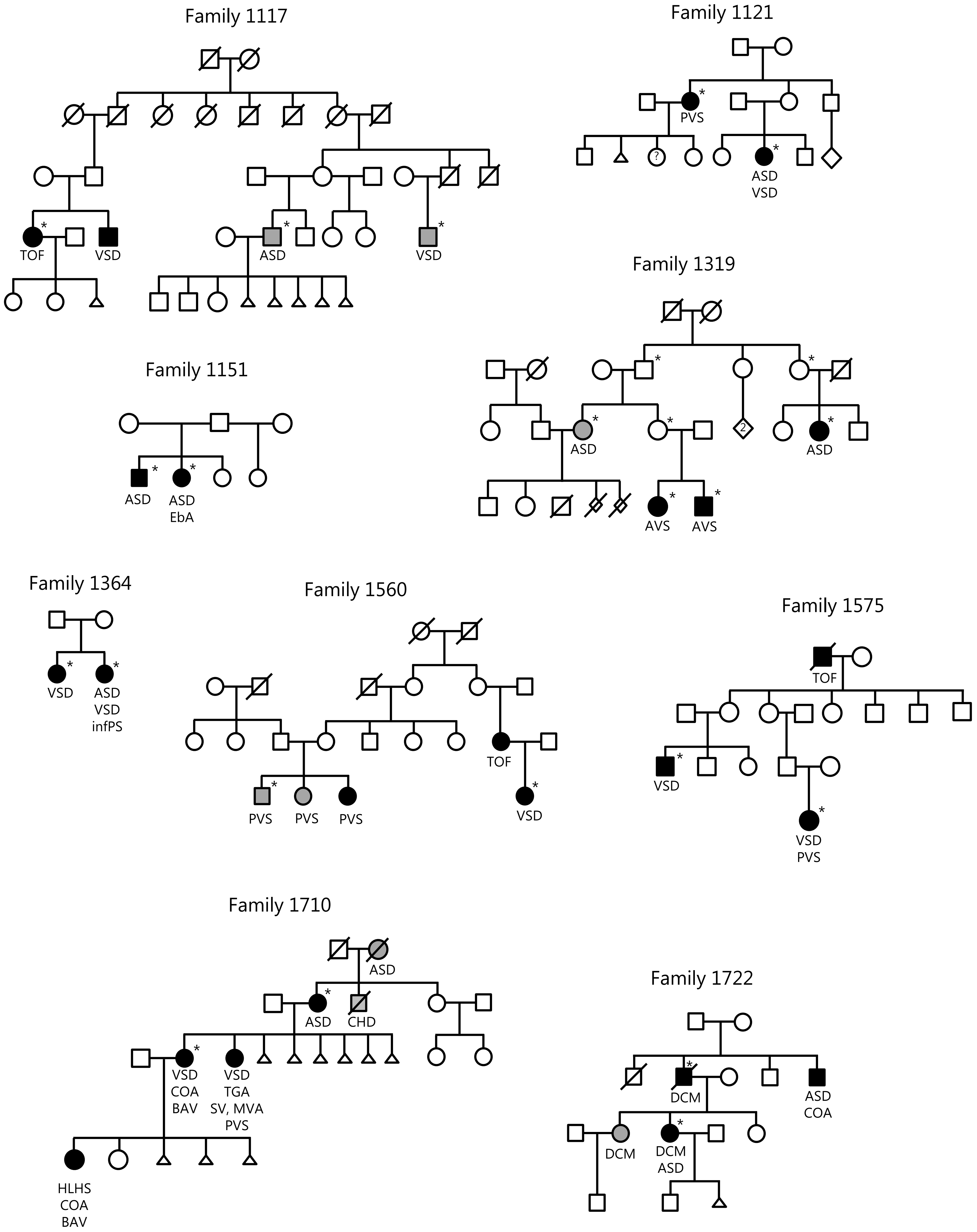
Family 1117
TOF
VSD
ASD
VSD
Family 1121
PVS
?
ASD
VSD
Family 1151
ASD
ASD
EbA
Family 1319
ASD
2
ASD
AVS
AVS
Family 1364
VSD
ASD
VSD
infPS
Family 1560
TOF
PVS
PVS
PVS
VSD
Family 1575
TOF
VSD
VSD
PVS
Family 1710
ASD
ASD
CHD
VSD
COA
BAV
VSD
TGA
SV, MVA
PVS
HLHS
COA
BAV
Family 1722
DCM
ASD
COA
DCM
DCM
ASD

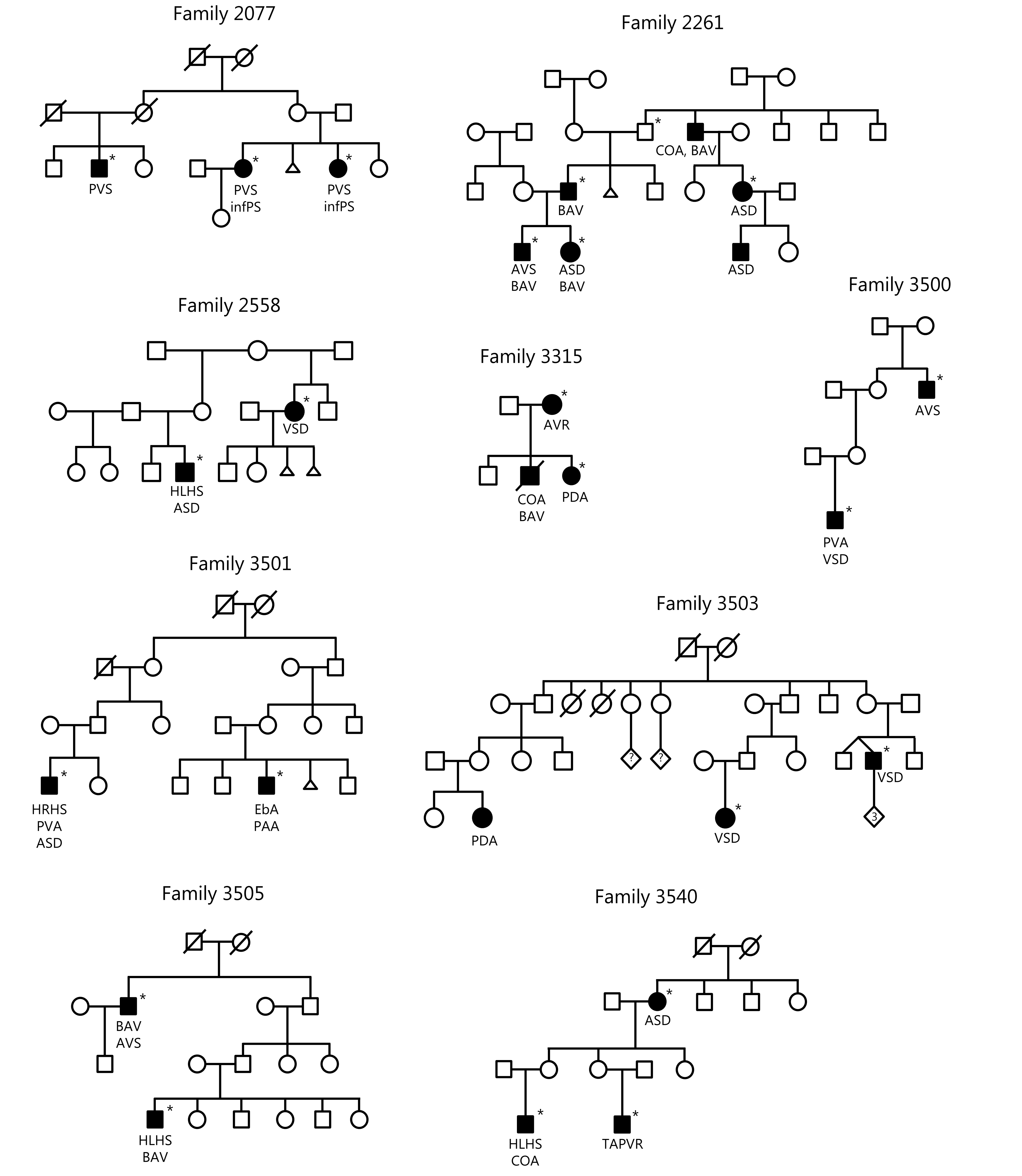

**Figure S1. Pedigrees of 32 Danish multiplex CHD families.** Circles: females. Squares: males. White symbols: unaffected family members. Filled symbols: affected family members. Triangles: abortion. CHD was determined by manual inspection of patient files (black) or data from DNPR/interview with family members (grey). Exome sequenced individuals are marked with asterix. The following heart defects were identified; Aortic Valve Regurgitation (AVR), Aortic Valve Stenosis (AVS), Atrial Septal Defect (ASD), Atrioventricular Septal Defect (AVSD), Bicuspid Aortic Valve(BAV), Coarctation of the Aorta (COA), Dilated Cardiomyopathy (DCM), Ebstein's Anomaly (EbA), Hypertrophic Cardiomyopathy (HCM), Hypoplastic Left Heart Syndrome (HLHS), Infundibular Pulmonary Stenosis (InfPS), Mitral Valve Atresia (MVA), Partial Anomalous Pulmonary Venous Return (PAPVR), Patent Ductus Arteriosus (PDA), Patent Foramen Ovale (PFO), Pulmonary Valve Atresia (PVA), Pulmonary Valve Stenosis (PVS), Single Ventricle (SV), Subvalvular Aortic Stenosis (SubAS), Tetralogy of Fallot (TOF), Total Anomalous Pulmonary Venous Return (TAPVR), Transposition of The Great Arteries (TGA), Vascular ring (Vring), Ventricular Septal Defect (VSD).

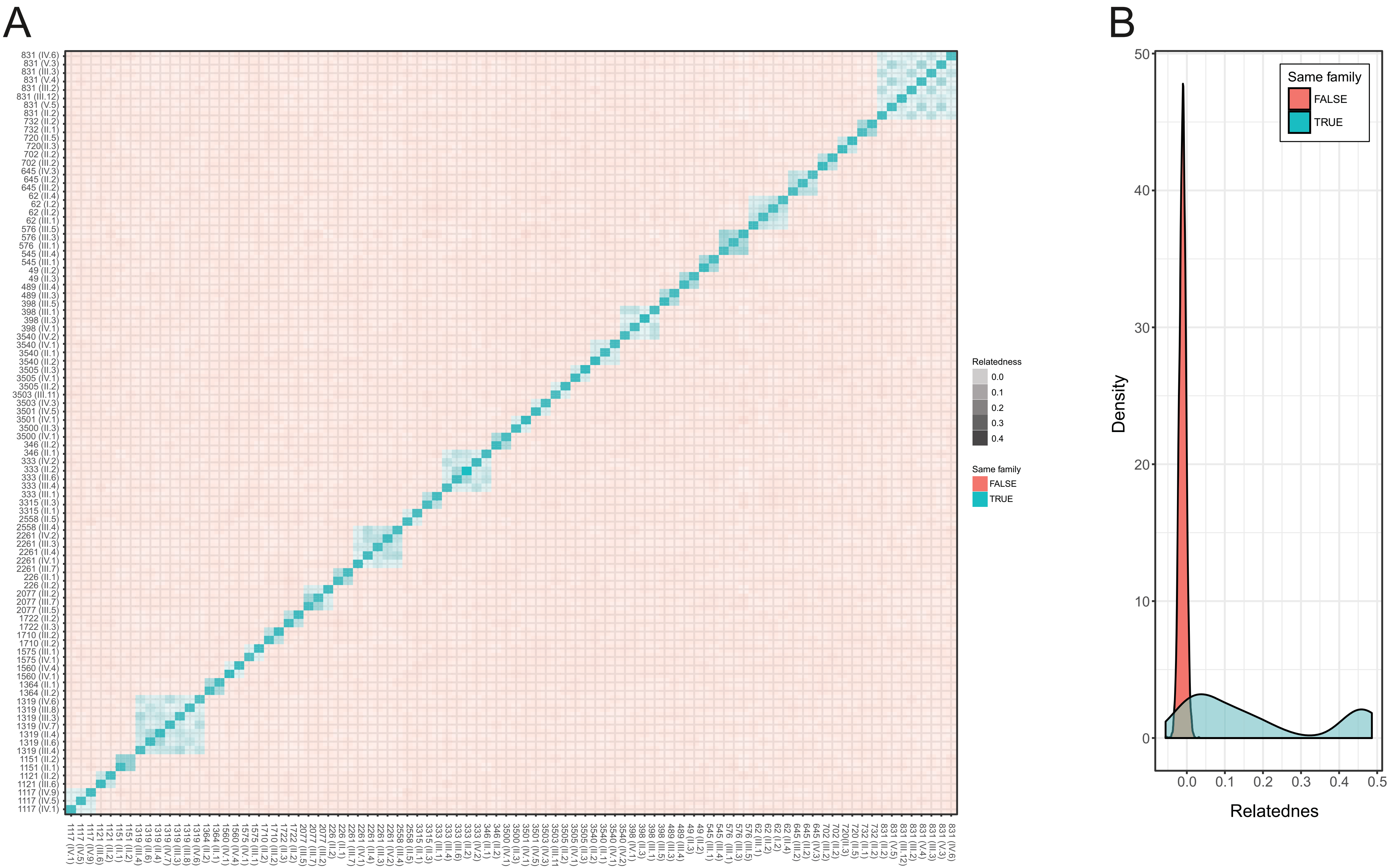

**Figure S2. Relatedness of the 90 individuals included in the study.** A. Heatmap showing the relatedness score of pairs of individuals. The family number and position in the family is indicated left and below the heatmap. B. Density plot of relatedness score from the pairwise analysis. Individuals from same family is shown in blue, and individuals not from the same family is shown in red.

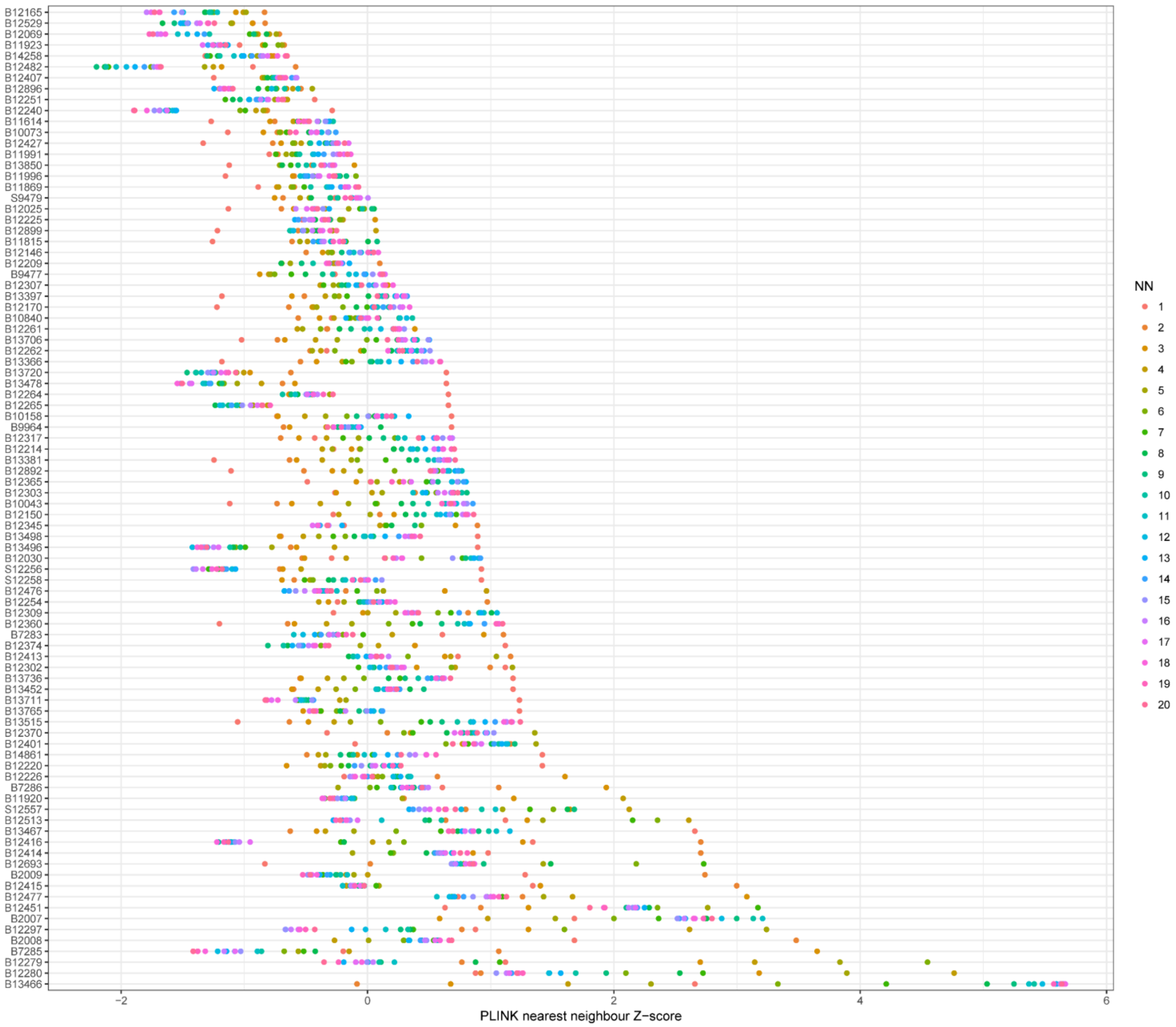

**Figure S3. Homogeneity of the cohort.** Population distances from each of the 90 individuals in the cohort to the 20 nearest neighbors (NN) were calculated. The distances compared to the the mean of the population in terms of standard deviations (Z-scores) is plotted on the x-axis.

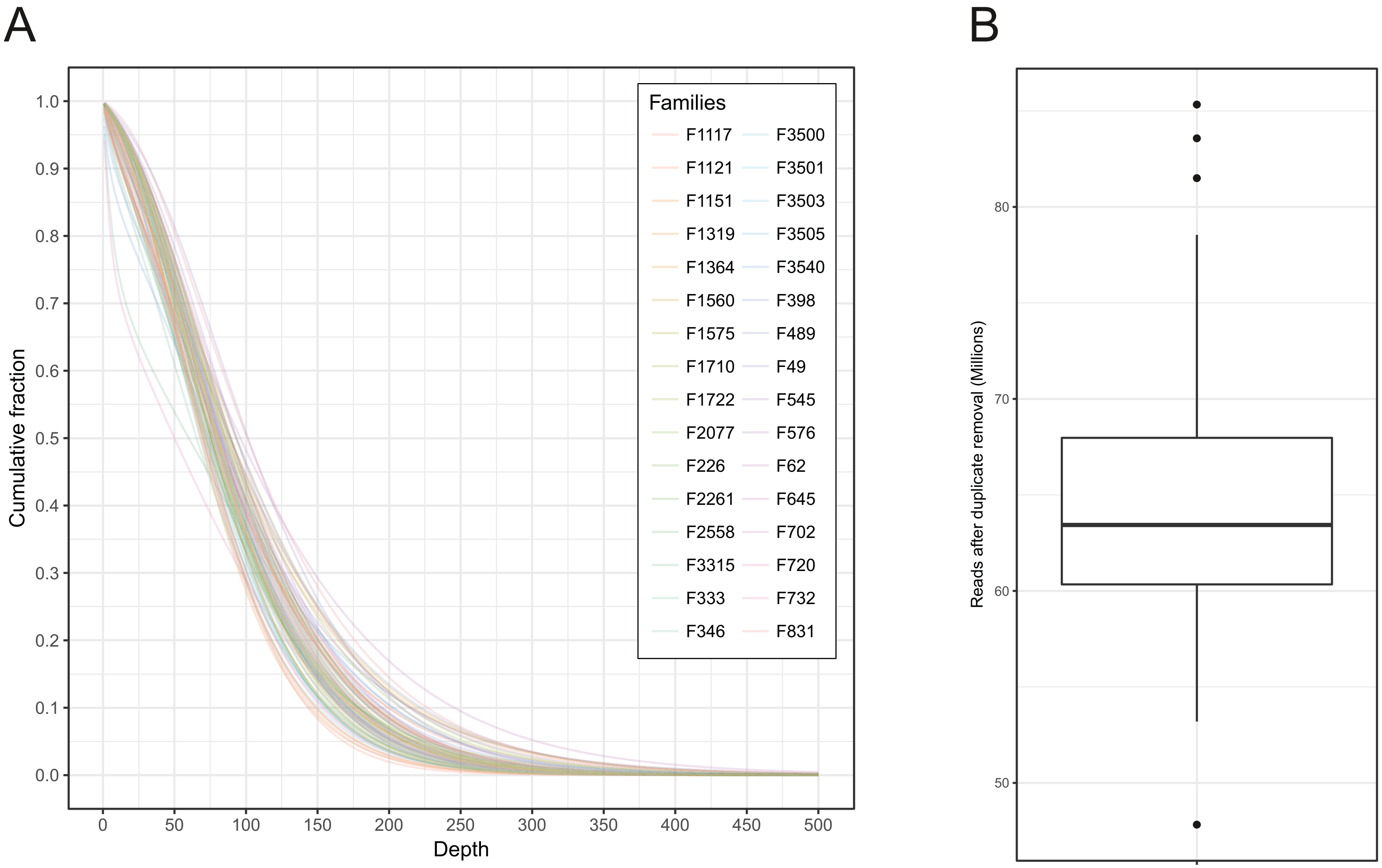

**Figure S4. Sequencing coverage and number of reads.** A. Cumulative depth of sequencing per family. B. Number of reads after removal of duplicates.

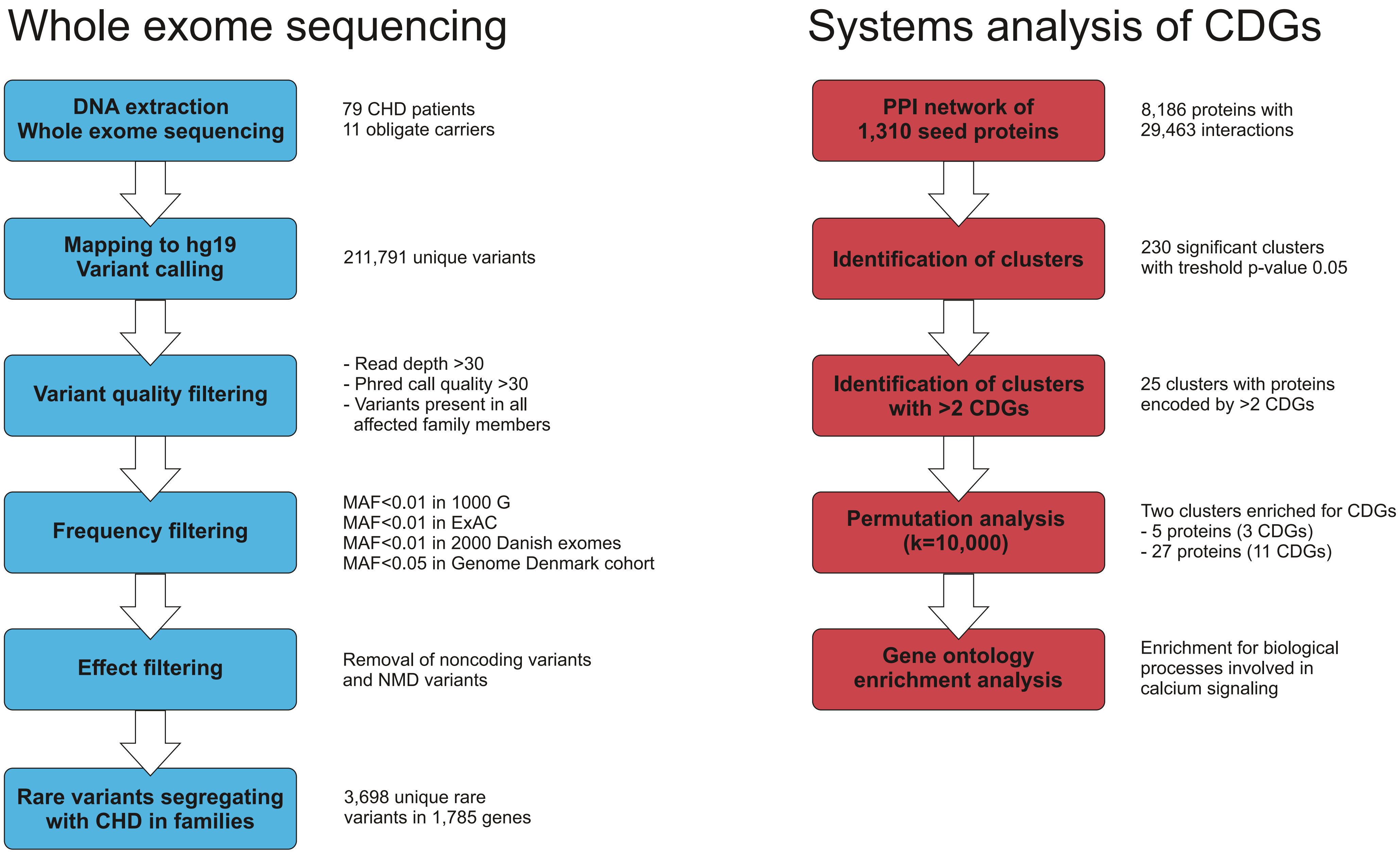

**Figure S5. Overview of the sequencing and data analysis processes.**

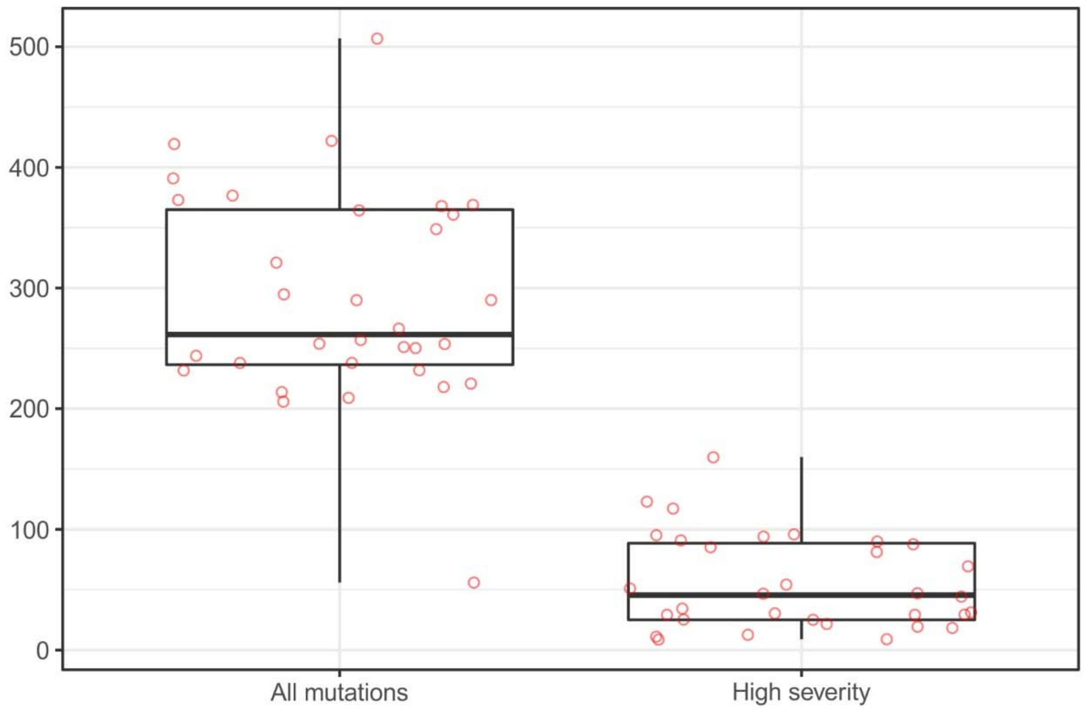

**Figure S6. Number of CDGs per family when all rare variants (left) or only high severity variants (right) were considered.** High severity variants were defined as variants creating splicing defects, frameshifts, premature stop codons and missense variants predicted to be damaging by both Polyphen and SIFT.

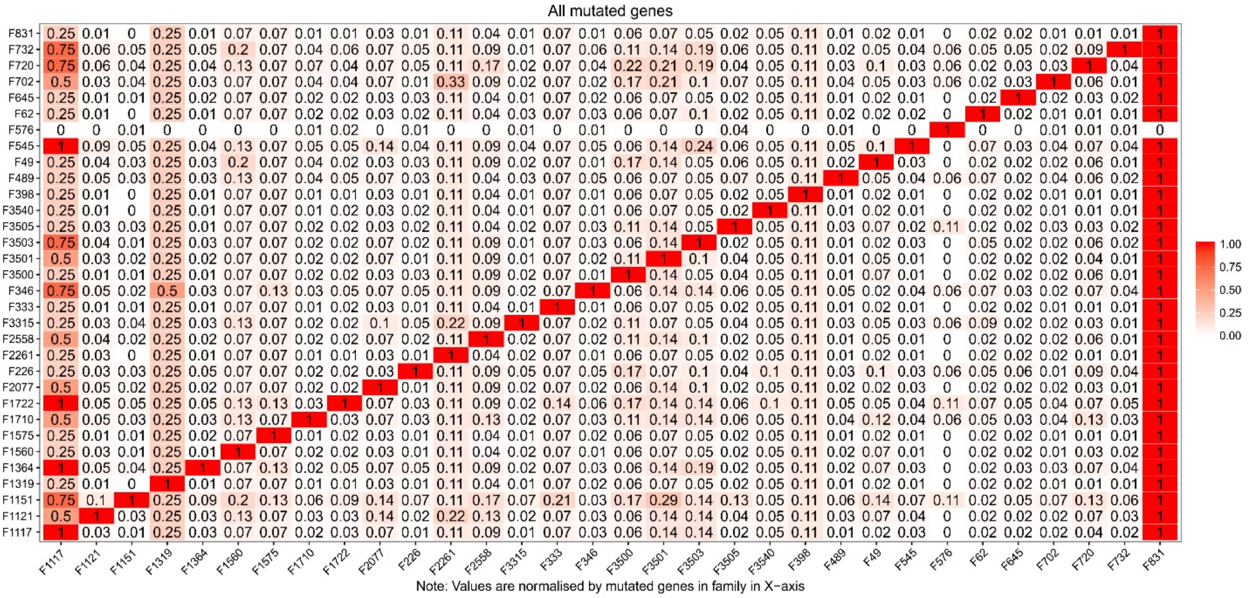

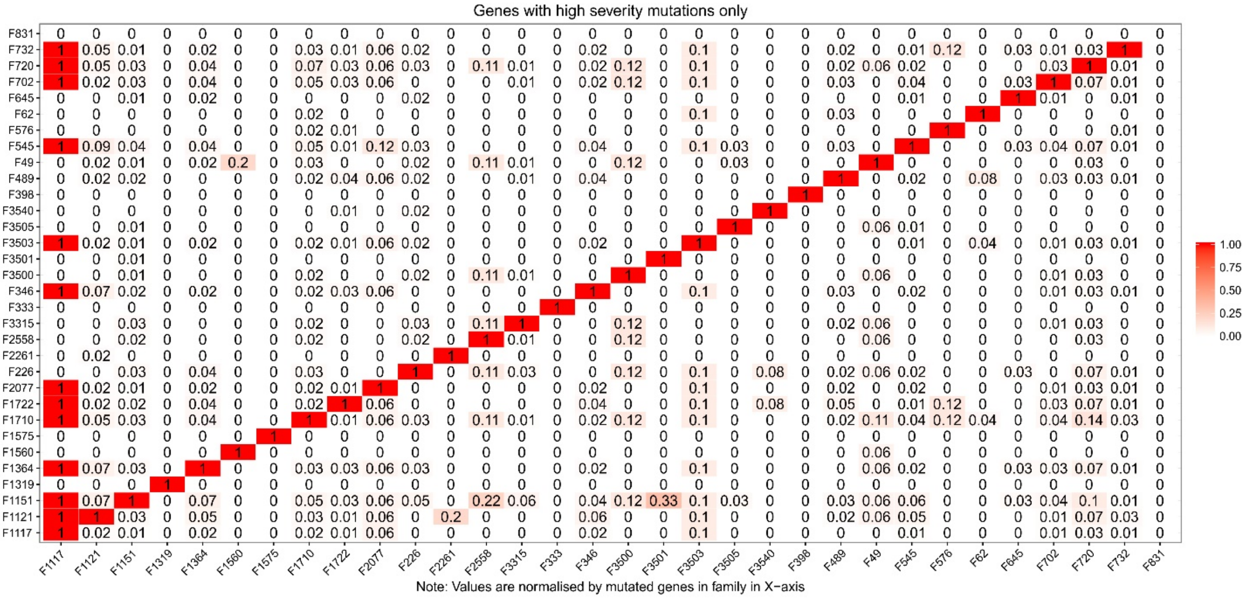

**Figure S7. Overlap between CDGs in pairs of families.** The fraction of overlap (FO) between CDGs are shown for each pair of families. For example, 11% of the CDGs identified in family 49 overlaps with CDGs identified in family 2558, which is indicated by a FO value of 0.11.

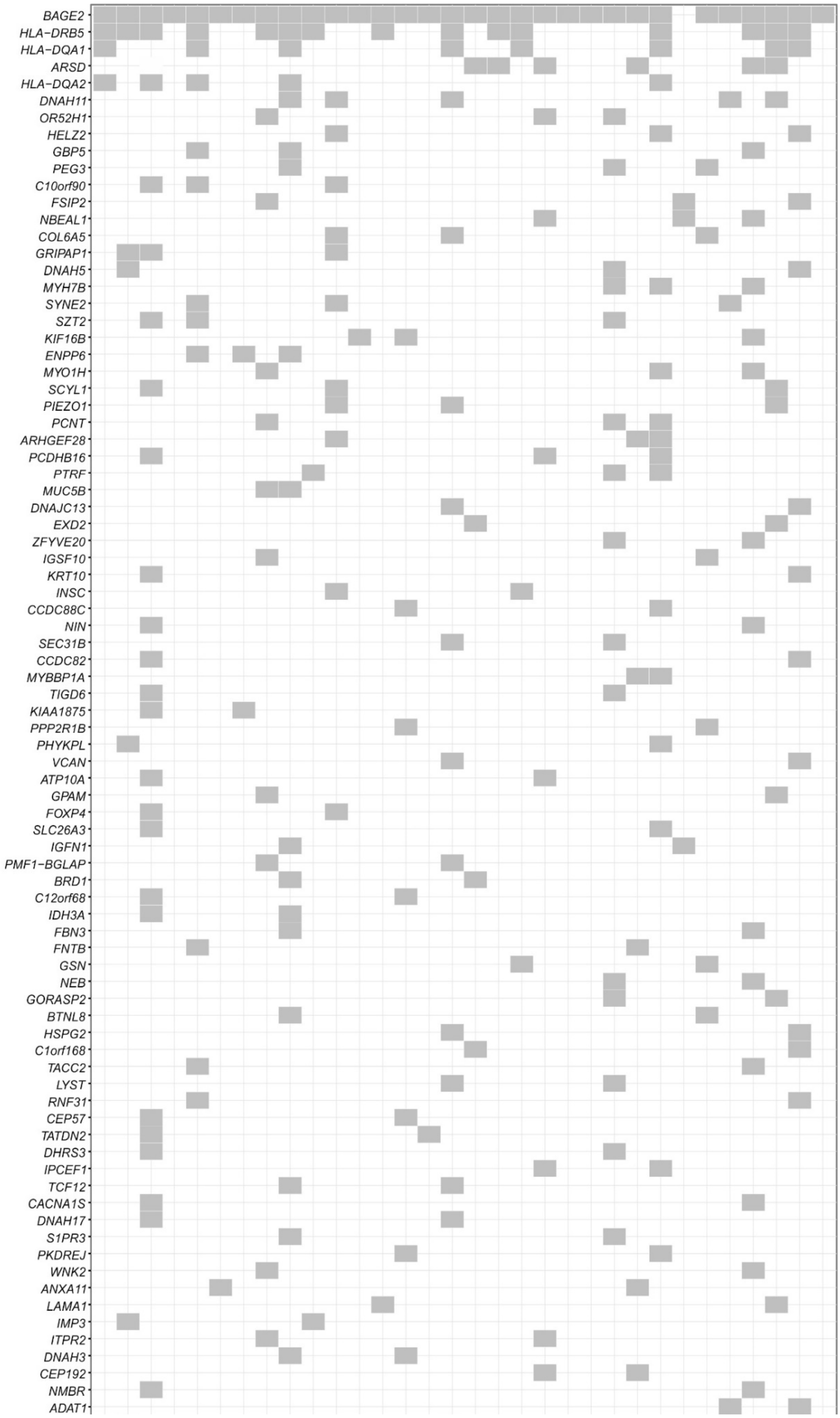
BAGE2
HLA−DRB5
HLA−DQA1
ARSD
HLA−DQA2
DNAH11
OR52H1
HELZ2
GBP5
PEG3
C10orf90
FSIP2
NBEAL1
COL6A5
GRIPAP1
DNAH5
MYH7B
SYNE2
SZT2
KIF16B
ENPP6
MYO1H
SCYL1
PIEZO1
PCNT
ARHGEF28
PCDHB16
PTRF
MUC5B
DNAJC13
EXD2
ZFYVE20
IGSF10
KRT10
INSC
CCDC88C
NIN
SEC31B
CCDC82
MYBBP1A
TIGD6
KIAA1875
PPP2R1B
PHYKPL
VCAN
ATP10A
GPAM
FOXP4
SLC26A3
IGFN1
PMF1−BGLAP
BRD1
C12orf68
IDH3A
FBN3
FNTB
GSN
NEB
GORASP2
BTNL8
HSPG2
C1orf168
TACC2
LYST
RNF31
CEP57
TATDN2
DHRS3
IPCEF1
TCF12
CACNA1S
DNAH17
S1PR3
PKDREJ
WNK2
ANXA11
LAMA1
IMP3
ITPR2
DNAH3
CEP192
NMBR
ADAT1

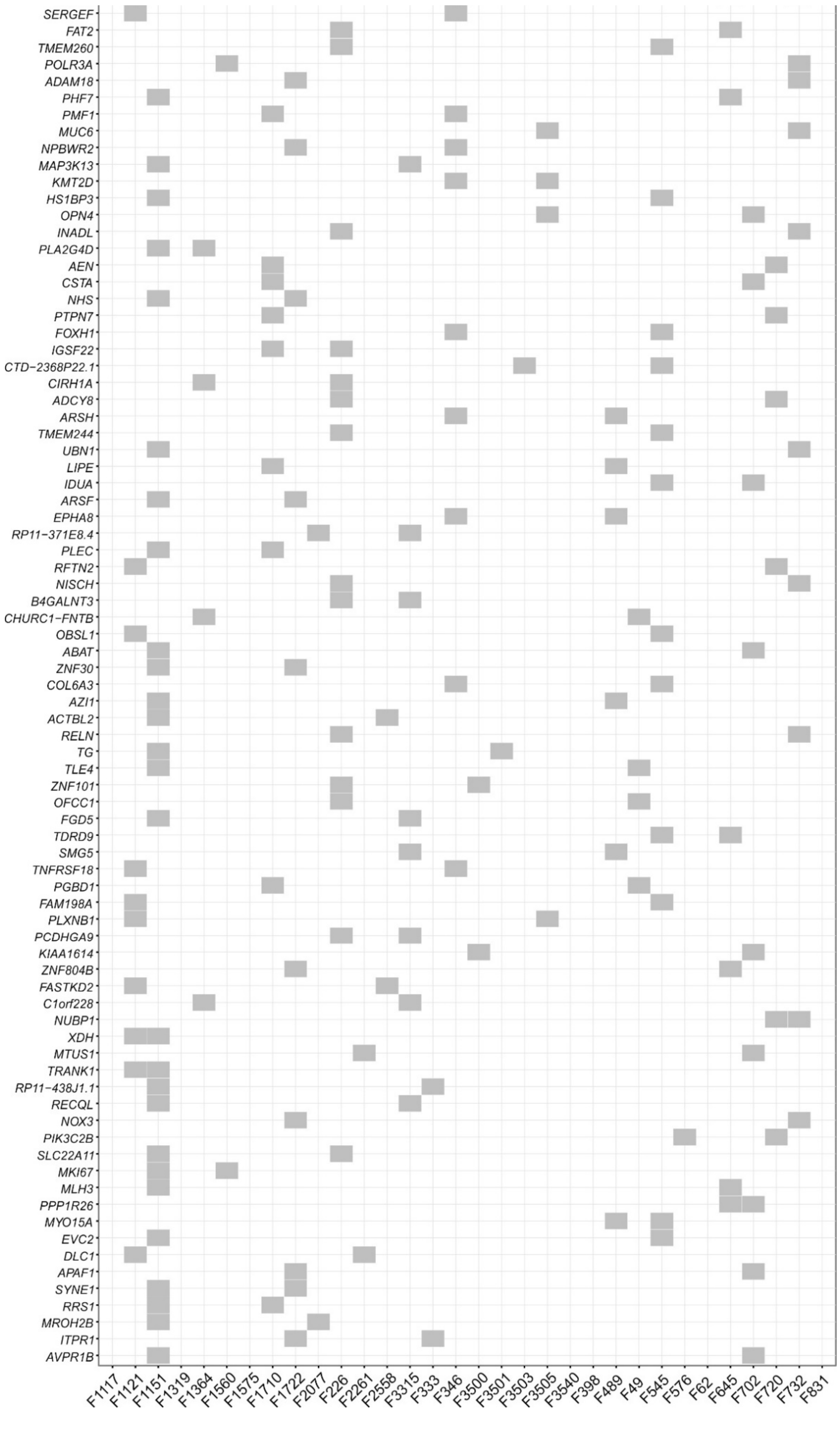

**Figure S8. Families with rare inherited variants in CDGs.** Only genes mutated in two or more families are shown.

a)

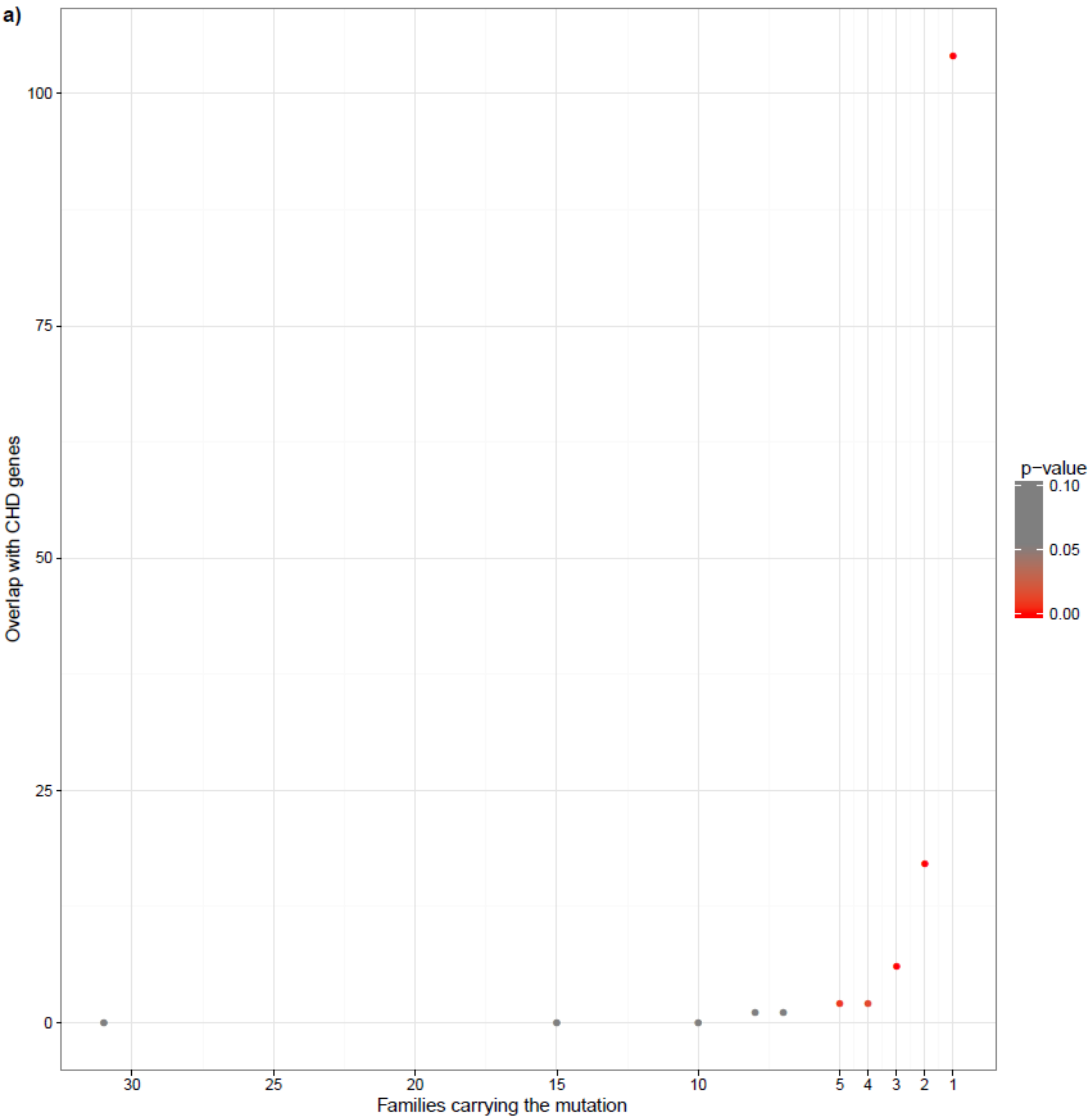
Overlap with CHD genes
100
75
50
25
0
30
25
20
15
10
5
4
3
2
1
Families carrying the mutation
p−value
0.10
0.05
0.00

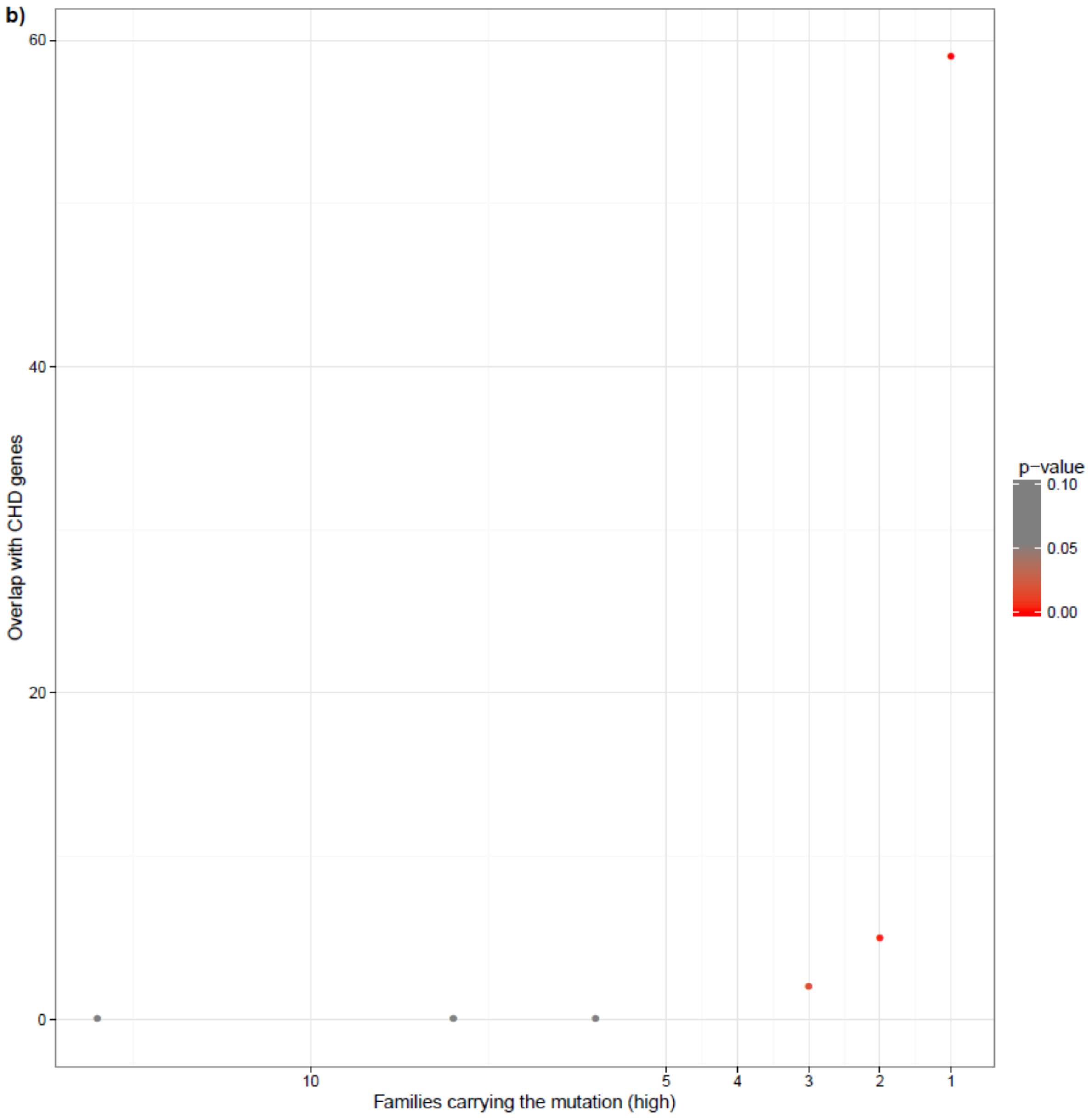

**Figure S9. Overlap between the 1,785 CDGs in our families and a curated list of 829 genes known to cause CHD in mice.** (A) Overlap with all 1,785 CDGs. (B) Overlap with genes carrying mutations scored pathogenic by SIFT/Polyphen-2.

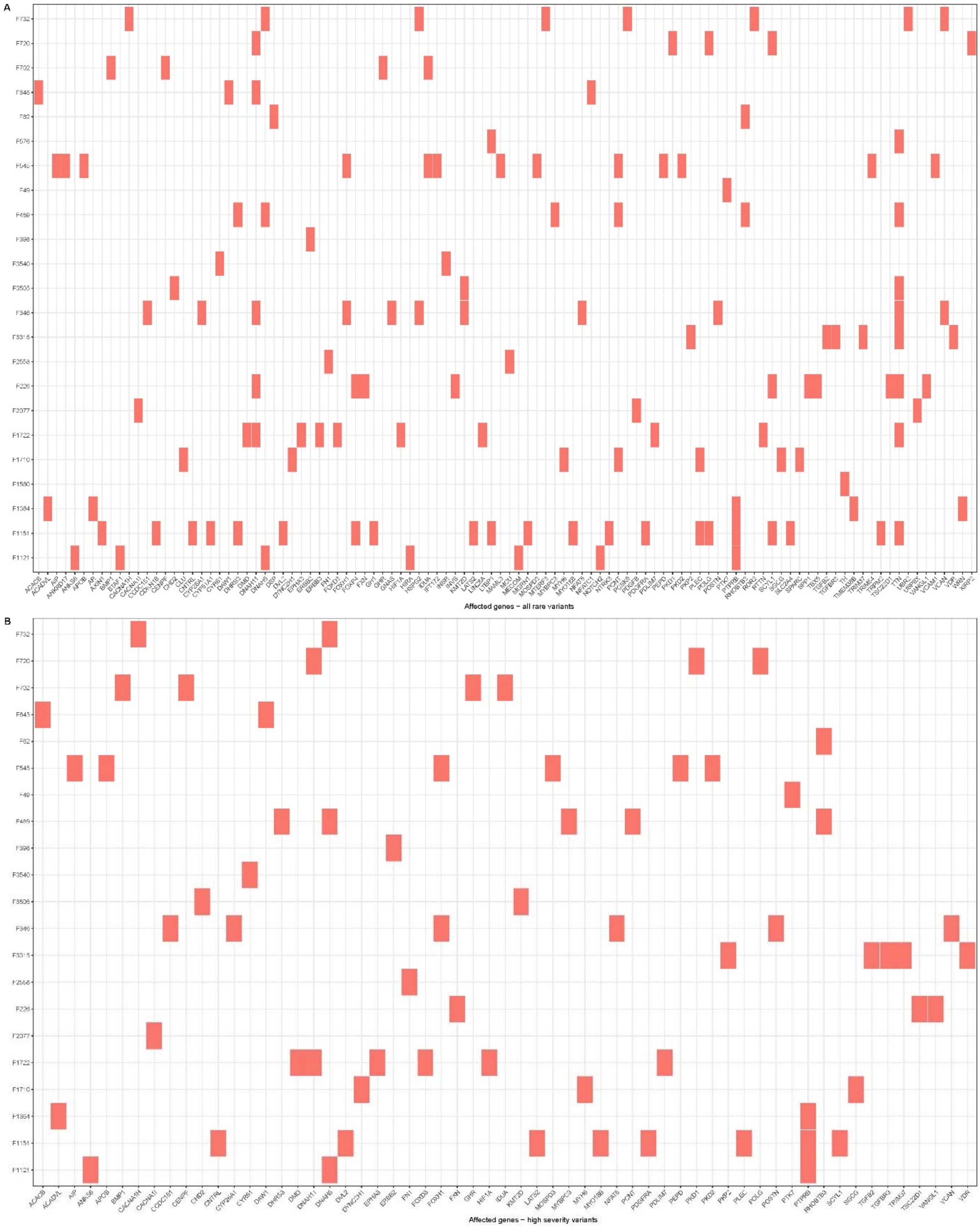

**Figure S10. Distribution of CHD genes across families.** CHD genes identified in mouse models are shown here. The distribution of human CHD genes is shown in Supplemental Table 4. (A) All CHD genes affected by rare variants. (B) CHD genes affected by high severity variants. High severity variants were defined as variants creating splicing defects, frameshifts, premature stop codons and missense variants predicted to be damaging by both Polyphen and SIFT.

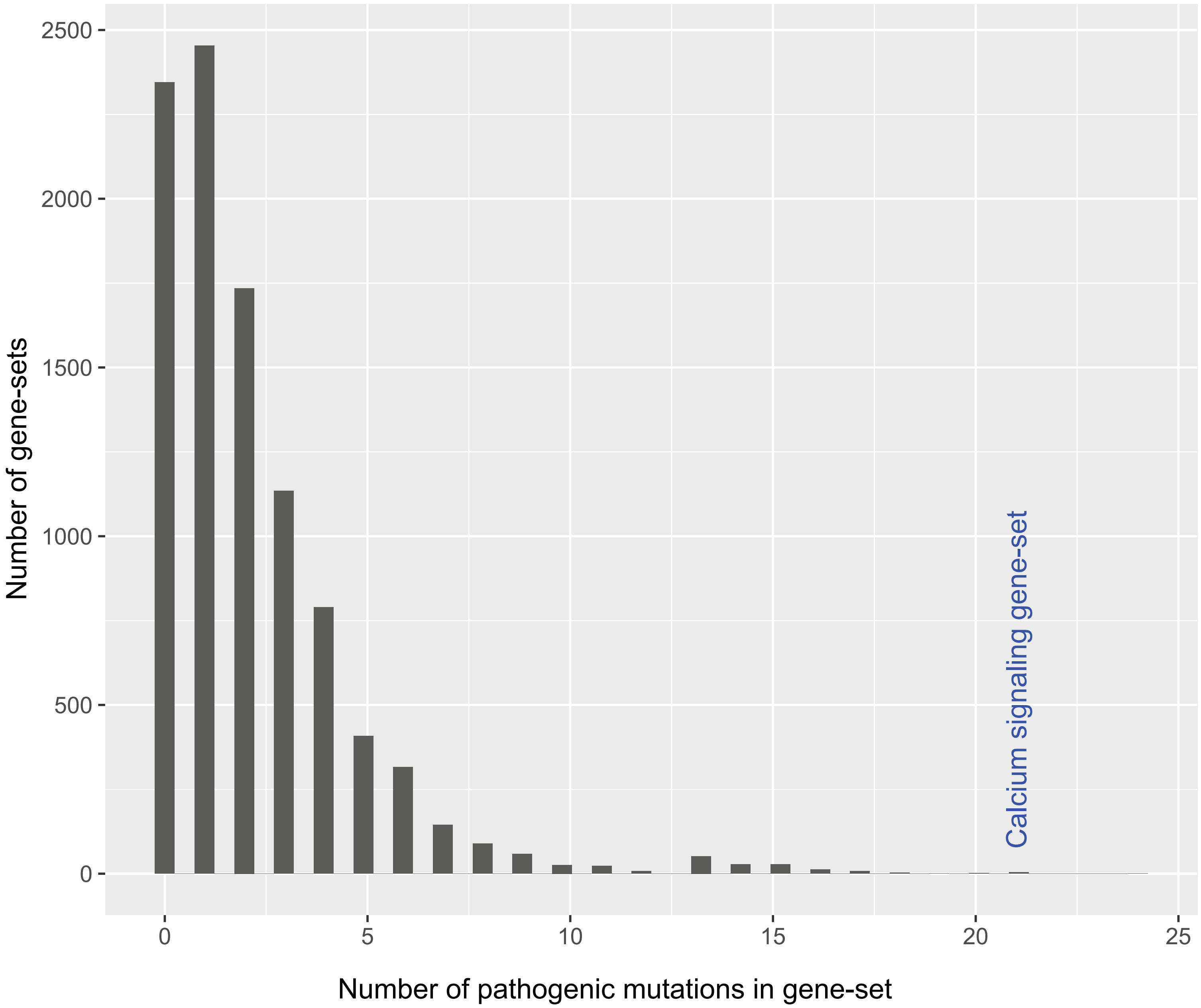

**Figure S11. Distribution of pathogenic mutations in random gene-sets.** 10,000 random gene-sets of 10 genes were created. The size distribution of each set was similar to the calcium-signaling gene-set tested in the case-control study; *ADCY2* (6575 bp), *ADCY5* (7311 bp), *ITPR1* (10197 bp), *CACNA1S* (6166 bp), *CACNA1I* (6740 bp), *CACNA1H* (8208 bp), *CACNA1D* (8991 bp), *GRIA4* (5508 bp), *PLCB2* (4616 bp), and *NFAT5* (13362 bp). Pathogenic mutations were defined as mutations with MAF < 0.001 and MPC score >2.

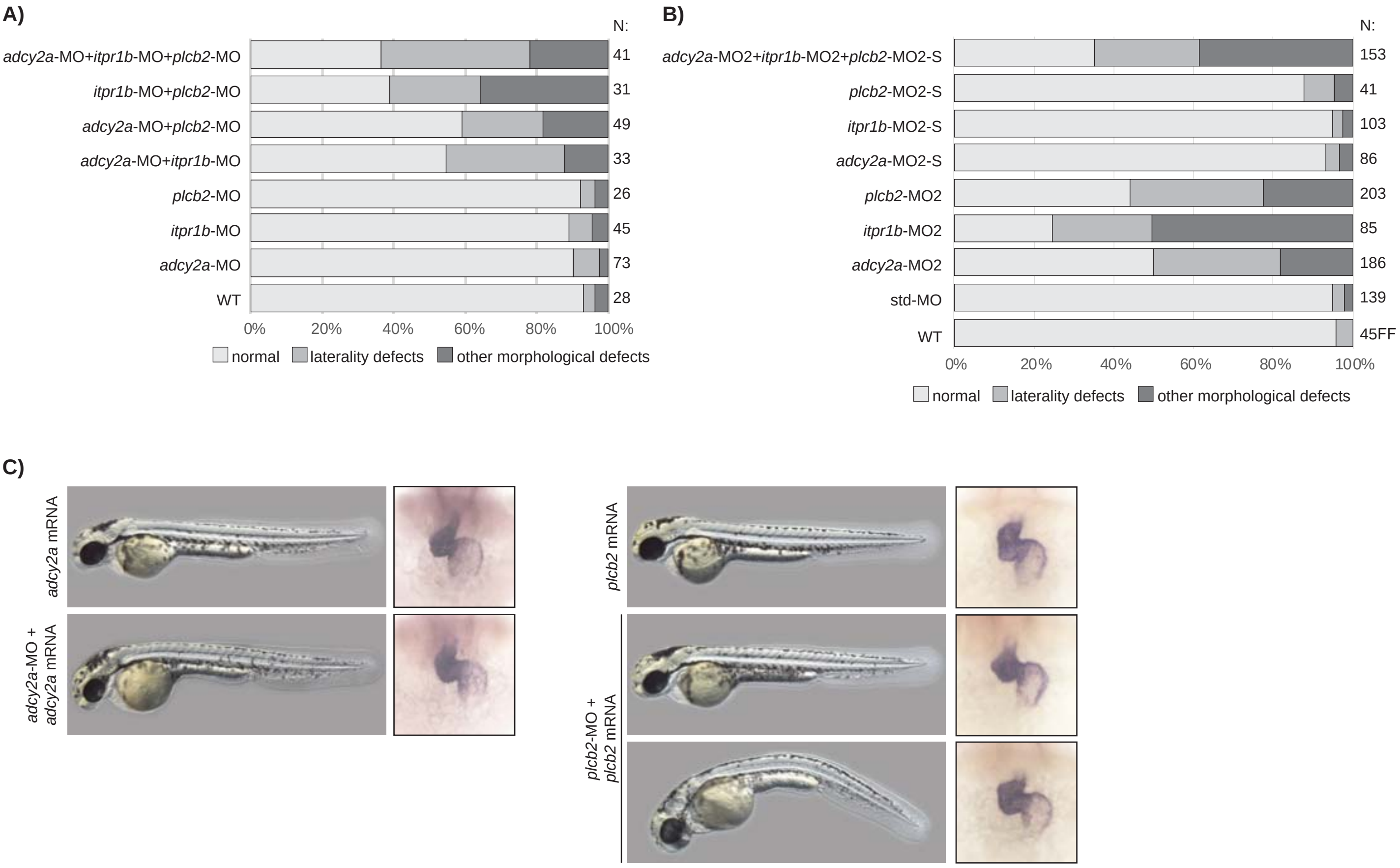

**Figure S12.** A. Injection of sub-efficient doses of MOs. Note that only combinations of MOs result in significant numbers of heart defects among the embryos. B. Quantification of heart phenotypes in WT, controls and morphants injected with second set of MOs. S indicates that sub-efficient doses of MOs are injected. C. Rescue of *adcy2a* and *plcb2* knockdown. Embryos injected with *adcy2a* or *plcb2* RNA alone and in combinations with corresponding MOs, are displayed at 48 hpf. *myl7* in situ hybridization marks the heart in these embryos.

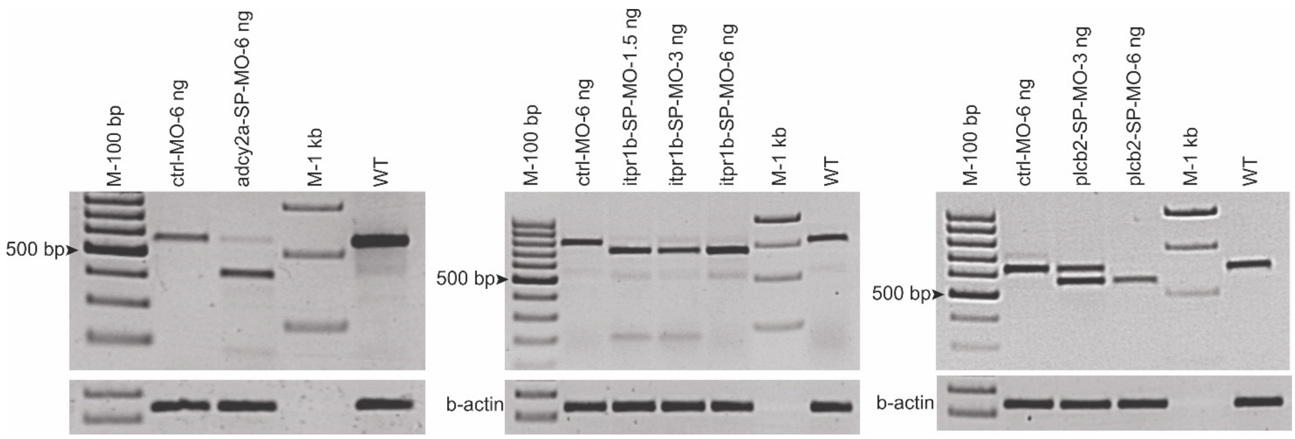

**Figure S13. Efficiency of the splice blocking morpholinos (MOs) used against *adcy2a*, *itpr1b* and *plcb2*.** Upper panel shows first set of MOs and lower panel shows second set of MOs. Aberrant splicing of *adcy2a*, *itpr1b* and *plcb2* and decrease in corresponding WT mature mRNA is confirmed by RT-PCR analysis.

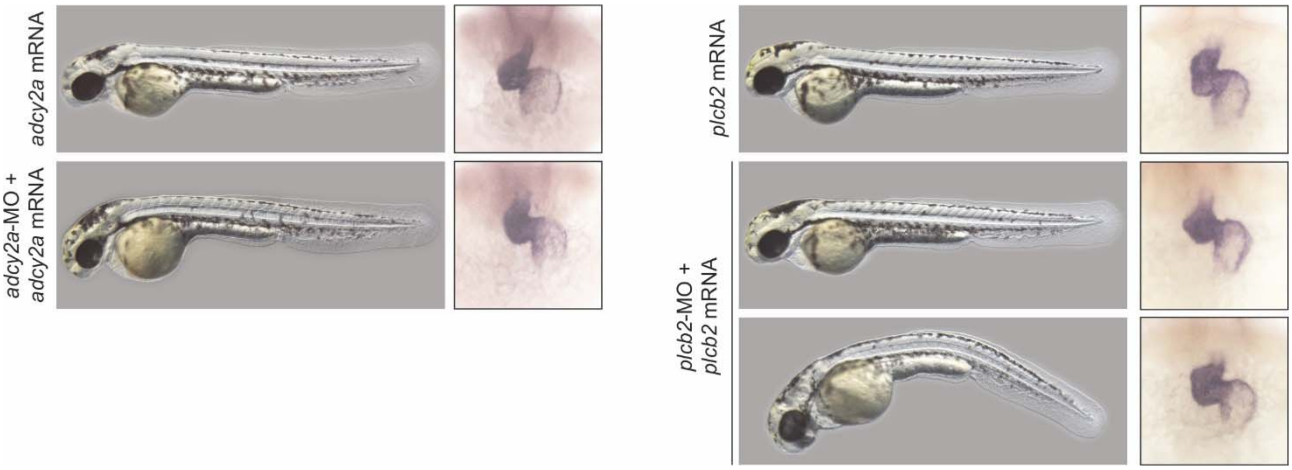

**Figure S14. Rescue of *adcy2a* and *plcb2* knockdown.** 48 hpf embryos injected with *adcy2a* or *plcb2* RNA alone and in combinations with corresponding SP-MOs, are displayed. *myl7* in situ hybridization marks the heart in these embryos.

**Table S1.** Human orthologues to 829 genes known to be associated with CHD in mouse models. The list was compiled from data in the Mouse Genome Informatics database (http://www.informatics.jax.org/).

| | | | | | | |
|---|---|---|---|---|---|---|
| *ABCA5* | *CHD7* | *FGFRL1* | *INVS* | *MTERF3* | *PPARG* | *SPTBN1* |
| *ABCB8* | *CHMP5* | *FHOD3* | *IRAK1* | *MTERF4* | *PPARGC1A* | *SRF* |
| *ABI1* | *CHRD* | *FKBP1A* | *IRX3* | *MTMR12* | *PPARGC1B* | *SRSF1* |
| *ABL1* | *CHST14* | *FKBP1B* | *IRX5* | *MTO1* | *PPP1R13L* | *SRSF10* |
| *ACACB* | *CISD2* | *FLNA* | *ISL1* | *MUS81* | *PPP2R3A* | *SSBP2* |
| *ACADM* | *CITED2* | *FLRT3* | *ITCH* | *MYBPC3* | *PPP2R5C* | *SSR1* |
| *ACADVL* | *CLU* | *FMOD* | *ITGA5* | *MYCN* | *PPP3CB* | *STK39* |
| *ACE* | *CLUAP1* | *FN1* | *ITGAV* | *MYH10* | *PRDM1* | *SUFU* |
| *ACKR3* | *CNTRL* | *FOXA2* | *ITGB1* | *MYH6* | *PRDM16* | *T* |
| *ACSL4* | *COL18A1* | *FOXC1* | *ITPA* | *MYH7* | *PRDM6* | *TAB1* |
| *ACTC1* | *COL1A1* | *FOXC2* | *JAG1* | *MYL2* | *PRF1* | *TAL1* |
| *ACVR1* | *COL4A3BP* | *FOXD3* | *JARID2* | *MYL7* | *PRICKLE1* | *TBC1D32* |
| *ACVR2A* | *COMMD9* | *FOXG1* | *JMJD6* | *MYLK3* | *PRKAR1A* | *TBX1* |
| *ACVR2B* | *CRB2* | *FOXH1* | *JPH2* | *MYO10* | *PRKCI* | *TBX18* |
| *ACVRL1* | *CREBBP* | *FOXJ1* | *JUN* | *MYO18B* | *PROC* | *TBX2* |
| *ADAM12* | *CRKL* | *FOXM1* | *JUND* | *MYOD1* | *PSEN1* | *TBX20* |
| *ADAM15* | *CSNK2A1* | *FOXO1* | *JUP* | *MYOZ2* | *PSEN2* | *TBX3* |
| *ADAM17* | *CSRP2* | *FOXP1* | *KAT6A* | *NACA* | *PSKH1* | *TBX5* |
| *ADAM19* | *CSRP3* | *FOXP4* | *KCNH2* | *NCKAP1* | *PTCD2* | *TBX6* |
| *ADAM9* | *CST9* | *FOXQ1* | *KCTD10* | *NCOA6* | *PTEN* | *TCF21* |
| *ADAMTS6* | *CTBP2* | *FRAS1* | *KDM1A* | *NCOR2* | *PTGS2* | *TCTN2* |
| *ADGRG6* | *CTNNA3* | *FREM2* | *KDM2A* | *NCSTN* | *PTK2* | *TDG* |
| *ADM* | *CTNNB1* | *FRS2* | *KDM4A* | *NDST1* | *PTK7* | *TEAD1* |
| *ADRA1A* | *CXADR* | *FSTL1* | *KDM6A* | *NEGR1* | *PTPN11* | *TEAD2* |
| *ADRA1B* | *CXCL12* | *FURIN* | *KDR* | *NEK6* | *PTPRB* | *TEK* |
| *AGT* | *CXCR4* | *FUS* | *KIF3A* | *NEK8* | *PTPRJ* | *TERC* |
| *AGTR1* | *CYBB* | *FUZ* | *KIF3B* | *NF1* | *RAC1* | *TFAP2A* |
| *AGTR2* | *CYBRD1* | *FXN* | *KIF7* | *NFAT5* | *RAF1* | *TFB1M* |
| *AIFM1* | *CYP11B2* | *FXR1* | *KIFAP3* | *NFATC1* | *RAMP2* | *TGFB1* |
| *AIP* | *CYP26A1* | *FZD1* | *KISS1R* | *NFATC3* | *RARA* | *TGFB2* |
| *AKAP12* | *CYP27B1* | *FZD2* | *KLF15* | *NFATC4* | *RARB* | *TGFBR2* |
| *AKAP13* | *CYP51A1* | *FZD7* | *KLF2* | *NGF* | *RARG* | *TGFBR3* |
| *AKAP6* | *CYR61* | *GAA* | *KLF3* | *NIPBL* | *RB1* | *TGIF1* |
| *AKT1* | *DAAM1* | *GAB1* | *KLF5* | *NKX2-5* | *RB1CC1* | *TGIF2* |
| *AKT3* | *DAG1* | *GAB2* | *KLHL40* | *NKX2-6* | *RBL2* | *TGM2* |
| *ALDH1A2* | *DAGLA* | *GAS1* | *KMT2B* | *NODAL* | *RBM20* | *TH* |
| *ALG13* | *DAND5* | *GATA4* | *KMT2D* | *NOG* | *RBP4* | *THBS1* |
| *ALPK3* | *DAW1* | *GATA5* | *KRAS* | *NOS1* | *RBPJ* | *THRA* |
| *AMN* | *DCHS1* | *GATA6* | *KRT19* | *NOS3* | *RCAN1* | *TIE1* |
| *ANGPT1* | *DCTN5* | *GBE1* | *LAMA4* | *NOTCH1* | *RCE1* | *TK2* |

| | | | | | | |
|---|---|---|---|---|---|---|
| *ANKRD17* | *DDR2* | *GBX2* | *LAMA5* | *NOTCH2* | *RDH10* | *TKT* |
| *ANKS6* | *DDX11* | *GDF1* | *LARGE1* | *NOV* | *RERE* | *TLL1* |
| *AP2B1* | *DDX3X* | *GH1* | *LATS2* | *NPPB* | *RFX3* | *TLR2* |
| *AP4E1* | *DEDD* | *GHR* | *LBX1* | *NPR1* | *RHBDF1* | *TMED2* |
| *APC* | *DES* | *GHRHR* | *LDB3* | *NPRL3* | *RHBDF2* | *TMEM100* |
| *APOB* | *DHRS3* | *GIPC1* | *LDLR* | *NR1D2* | *RHEB* | *TMEM106B* |
| *APOE* | *DICER1* | *GJA1* | *LEFTY1* | *NR1H3* | *RHOBTB3* | *TMEM38A* |
| *AR* | *DISP1* | *GJA5* | *LEFTY2* | *NR2F2* | *RIC8B* | *TMEM38B* |
| *ARID1A* | *DLL1* | *GJC1* | *LEMD2* | *NR3C1* | *RIPPLY3* | *TMEM67* |
| *ARID2* | *DLL4* | *GNA11* | *LEP* | *NR3C2* | *RNF4* | *TMOD1* |
| *ARMC4* | *DMD* | *GNAQ* | *LEPR* | *NRG1* | *ROBO1* | *TMSB4X* |
| *ARNTL* | *DNAAF2* | *GNAS* | *LHCGR* | *NRP1* | *ROR1* | *TNNI3* |
| *ARSB* | *DNAAF3* | *GNG5* | *LHX1* | *NRP2* | *ROR2* | *TNNT2* |
| *ASL* | *DNAH11* | *GPC3* | *LIG3* | *NTF3* | *RPGRIP1L* | *TP53* |
| *ATE1* | *DNAH5* | *GRIN2D* | *LIMS1* | *NTRK3* | *RPS6KA2* | *TREX1* |
| *ATF2* | *DNAI1* | *GRK2* | *LIMS2* | *NXN* | *RSPO3* | *TRIM37* |
| *ATF7* | *DNAJA3* | *GSC* | *LIN28A* | *OFD1* | *RTTN* | *TRIM54* |
| *ATG5* | *DNASE1L2* | *GTF2I* | *LMNA* | *OPA3* | *RXRA* | *TRIM55* |
| *ATMIN* | *DNM1L* | *GTF2IRD1* | *LMOD2* | *OSR1* | *RXRB* | *TRIM63* |
| *ATP2A2* | *DNM2* | *GYS1* | *LOXL2* | *OTX2* | *RYR1* | *TRIP11* |
| *AXIN1* | *DNMT3B* | *GYS2* | *LRP1* | *OVOL2* | *RYR2* | *TRPM2* |
| *AXIN2* | *DOCK1* | *HADHB* | *LRP2* | *PAK1* | *SALL1* | *TSC1* |
| *B9D1* | *DOT1L* | *HAND1* | *LTBP1* | *PAK4* | *SALL4* | *TSC22D1* |
| *BAG3* | *DRC1* | *HAND2* | *LTBP4* | *PARD3* | *SCN5A* | *TTBK2* |
| *BAZ1B* | *DSP* | *HAS2* | *LUM* | *PARVA* | *SCN8A* | *TTC7A* |
| *BBX* | *DTNA* | *HBEGF* | *LUZP1* | *PATZ1* | *SCYL1* | *TTN* |
| *BCAR1* | *DUSP8* | *HCCS* | *LY6E* | *PAX3* | *SEC24B* | *TXNRD2* |
| *BCL6* | *DVL1* | *HDAC2* | *MAML1* | *PAXIP1* | *SEMA3C* | *UBE2U* |
| *BCOR* | *DVL2* | *HDAC5* | *MAML3* | *PBRM1* | *SEMA3D* | *UBE4B* |
| *BICC1* | *DVL3* | *HDAC9* | *MAP1S* | *PCNT* | *SENP2* | *UBP1* |
| *BIN1* | *DYNC2H1* | *HECTD1* | *MAP2K1* | *PCSK5* | *SGCB* | *UBR1* |
| *BMP1* | *DYNC2LI1* | *HECTD3* | *MAP2K2* | *PCSK6* | *SGCD* | *UBR2* |
| *BMP10* | *DYNLL1* | *HEG1* | *MAP2K5* | *PDCD10* | *SGCG* | *UBR4* |
| *BMP2* | *DYRK1A* | *HERC4* | *MAP3K3* | *PDGFA* | *SGPL1* | *USP12* |
| *BMP4* | *DYX1C1* | *HEXIM1* | *MAP3K7* | *PDGFB* | *SH3PXD2B* | *USP24* |
| *BMPR1A* | *ECE1* | *HEY1* | *MAPK1* | *PDGFC* | *SHC1* | *USP9X* |
| *BMPR2* | *ECE2* | *HEY2* | *MAPK11* | *PDGFRA* | *SHH* | *UTF1* |
| *BRAF* | *EDN1* | *HEYL* | *MAPK14* | *PDGFRB* | *SHOC2* | *UTRN* |
| *BTAF1* | *EDNRA* | *HGS* | *MAPK3* | *PDHA1* | *SHOX2* | *UVRAG* |
| *BTC* | *EDNRB* | *HHEX* | *MAPK6* | *PDLIM3* | *SIRT1* | *VANGL1* |
| *C2CD3* | *EFNA1* | *HIC2* | *MAPK7* | *PDLIM7* | *SIRT7* | *VANGL2* |
| *CACNA1C* | *EFNB2* | *HIF1A* | *MARVELD2* | *PDPK1* | *SIX1* | *VAV2* |
| *CACNA1H* | *EGFR* | *HIF3A* | *MATR3* | *PDPN* | *SLC22A8* | *VAV3* |

| | | | | | | |
|---|---|---|---|---|---|---|
| *CACNA1I* | *EGLN1* | *HIRA* | *MB* | *PDS5A* | *SLC2A4* | *VCAM1* |
| *CACNB2* | *EGLN3* | *HMOX1* | *MBTD1* | *PDS5B* | *SLC38A10* | *VCAN* |
| *CALCRL* | *EGR2* | *HOPX* | *MCU* | *PELI1* | *SLC39A4* | *VCL* |
| *CALR* | *EHMT1* | *HOXA1* | *MDM2* | *PEPD* | *SLC4A1* | *VDR* |
| *CAPNS1* | *EIF2B5* | *HOXA3* | *MECOM* | *PHC1* | *SLC6A6* | *VEGFA* |
| *CASP8* | *ELN* | *HOXB4* | *MED1* | *PHIP* | *SLC8A1* | *VEGFB* |
| *CASQ2* | *EMC10* | *HPRT1* | *MED12* | *PIBF1* | *SLC9A1* | *VPS54* |
| *CAV1* | *EMD* | *HRAS* | *MED24* | *PIFO* | *SLIT3* | *WASF2* |
| *CAV3* | *ENG* | *HSD11B2* | *MED30* | *PIGV* | *SMAD1* | *WDPCP* |
| *CC2D2A* | *EP300* | *HSD17B7* | *MEF2C* | *PIK3CA* | *SMAD2* | *WDR1* |
| *CCDC124* | *EPHA3* | *HSPB11* | *MEF2D* | *PIK3CB* | *SMAD3* | *WDR35* |
| *CCDC151* | *EPHB4* | *HSPB8* | *MEGF8* | *PIK3CG* | *SMAD4* | *WHSC1* |
| *CCDC160* | *EPO* | *HSPG2* | *MEIS1* | *PIK3R1* | *SMAD5* | *WNK1* |
| *CCDC39* | *EPOR* | *HTR2B* | *MEN1* | *PIK3R2* | *SMAD6* | *WNT11* |
| *CCM2* | *ERBB2* | *HTRA2* | *MEOX2* | *PIKFYVE* | *SMAD7* | *WNT3A* |
| *CCM2L* | *ERBB3* | *HUWE1* | *MESP1* | *PINK1* | *SMARCA4* | *WNT5A* |
| *CCND1* | *ERBB4* | *IDUA* | *MEST* | *PIP5K1C* | *SMG1* | *WRN* |
| *CCND2* | *ESR1* | *IER3* | *MFN2* | *PITX2* | *SMG9* | *WT1* |
| *CCND3* | *ESR2* | *IFT122* | *MFSD8* | *PKD1* | *SMN1* | *WWTR1* |
| *CCNE1* | *EYA1* | *IFT140* | *MGAT1* | *PKD2* | *SMO* | *XBP1* |
| *CCNE2* | *EZH2* | *IFT172* | *MGP* | *PKP2* | *SMYD1* | *XIRP1* |
| *CDC73* | *F11* | *IFT27* | *MGRN1* | *PLA2G4A* | *SNAI1* | *XIRP2* |
| *CDH2* | *F2R* | *IFT46* | *MIB1* | *PLAGL1* | *SNAI2* | *YAP1* |
| *CDH5* | *FADD* | *IFT57* | *MIXL1* | *PLCE1* | *SNX17* | *YWHAE* |
| *CDK2* | *FAT4* | *IFT74* | *MKL1* | *PLEC* | *SNX27* | *ZBTB14* |
| *CDK4* | *FBLN1* | *IFT88* | *MKL2* | *PLN* | *SOD1* | *ZFP36L1* |
| *CDK6* | *FBN1* | *IGF1R* | *MKS1* | *PLVAP* | *SOD2* | *ZFPM1* |
| *CDKN1A* | *FENDRR* | *IGF2* | *MMP21* | *PLXND1* | *SOS1* | *ZFPM2* |
| *CDKN1B* | *FES* | *IGF2R* | *MMP9* | *PNN* | *SOX11* | *ZIC3* |
| *CENPF* | *FGF10* | *IGFBP2* | *MORF4L1* | *PNPLA2* | *SOX12* | *ZNF366* |
| *CENPJ* | *FGF16* | *IGHMBP2* | *MOSPD3* | *POFUT1* | *SOX4* | *ZNF521* |
| *CEP290* | *FGF2* | *IKBKAP* | *MPZL3* | *POGLUT1* | *SOX9* | *ZSCAN10* |
| *CFC1* | *FGF8* | *IL17RD* | *MSX1* | *POLG* | *SPARC* | |
| *CFL1* | *FGF9* | *IL6* | *MSX2* | *POR* | *SPP1* | |
| *CFLAR* | *FGFR1* | *INSR* | *MT-CO1* | *POSTN* | *SPTA1* | |
| *CHD2* | *FGFR2* | *INTU* | *MT-RNR2* | *PPARA* | *SPTAN1* | |

**Table S2.** A list of 144 Human CHD disease genes. The list was curated from the literature (PMIDs 29106500, 23934094, 27058611, 28991257, 27479907, 28592524, 25996639, 29089047).

| | | | | | | |
|---|---|---|---|---|---|---|
| *ACTC1* | *CITED2* | *FLT4* | *KIAA0196* | *MYH7* | *PKD2* | *SOS1* |
| *ACVR1* | *COL2A1* | *FOXC1* | *KMT2A* | *NF1* | *PLRG1* | *SRCAP* |
| *ACVR2B* | *COL9A1* | *FOXC2* | *KMT2D* | *NFATC1* | *PRDM1* | *STK4* |
| *ADNP* | *CREBBP* | *FOXH1* | *KRAS* | *NIPBL* | *PRKD1* | *STRA6* |
| *ALDH1A2* | *CRELD1* | *FOXL2* | *LBR* | *NKX2* | *PTGFRA* | *TAB2* |
| *ALK2* | *CYR61* | *FOXP1* | *LEFTY2* | *NKX2-5* | *PTPN11* | *TBX1* |
| *ANKRD1* | *DDX3X* | *GATA4* | *MAP2K1* | *NKX2-6* | *RAF1* | *TBX20* |
| *ANKRD11* | *DHCR7* | *GATA5* | *MAP2K2* | *NODAL* | *RBFOX2* | *TBX3* |
| *B3GALTL* | *DNAH11* | *GATA6* | *MCTP2* | *NOTCH1* | *RBM10* | *TBX5* |
| *B3GAT3* | *DNAH5* | *GDF1* | *MDM4* | *NOTCH2* | *RIT1* | *TDGF1* |
| *BBS2* | *DNAI1* | *GDF3* | *MED13L* | *NPHP2* | *RNF20* | *TFAP2B* |
| *BCOR* | *DVL1* | *GJA5* | *MEK1* | *NPHP3* | *ROBO1* | *TNNI3* |
| *BMPR1A* | *DYRK1A* | *GPC3* | *MESP1* | *NPHP4* | *ROR2* | *VCAN* |
| *BRAF* | *EHMT1* | *HAND2* | *MID1* | *NR1D2* | *SALL1* | *WNT5A* |
| *CBL* | *ELN* | *HAS2* | *MKKS* | *NR2F2* | *SALL4* | *ZEB2* |
| *CCDC11* | *EP300* | *HOXA1* | *MKRN2* | *NRAS* | *SEMA3D* | *ZFMP2* |
| *CDK13* | *EVC* | *HRAS* | *MKS1* | *NRP1* | *SH3PXD2B* | *ZFPM2* |
| *CEP152* | *EVC1* | *IRX4* | *MLL2* | *NSD1* | *SHOC2* | *ZIC3* |
| *CFC1* | *EVC2* | *JAG1* | *MMP21* | *PACS1* | *SHROOM3* | |
| *CHD4* | *FBN1* | *JARID2* | *MYH11* | *PDGFRA* | *SMAD6* | |
| *CHD7* | *FGFR3* | *KDM6A* | *MYH6* | *PITX2* | *SMURF1* | |

**Table S3.** Primers used in the study

| Name | 5´-3´sequence |
|---|---|
| *Primers for test of morpholinos* | |
| zAdcy2a-SP-MO-test-F | GGAGTTTGAAAAGCGTCAGC |
| zAdcy2a-SP-MO-test-R | GGTACACCACCTGCTTCCAT |
| zItpr1b-SP-MO-test-F | TGTGGAGGTGGTGAGAAAGC |
| zItpr1b-SP-MO-test-R | AGAGGCTGTCCTGCGTTTAC |
| zPlcb2-SP-MO-test-F | ACGCTCTGCTTATCGACCTG |
| zPlcb2-SP-MO-test-R | GTCTGGTAGTTGAGGGCCAC |
| zAdcy2a-SP-MO2-test-F | GCCAGTCTGCAGTTCAACAT |
| zAdcy2a-SP-MO2-test-R | TGCCAGAATCTGCCATACCA |
| zPlcb2-SP-MO2-test-F | CTCTGATGAGGGAACGGCTG |
| zPlcb2-SP-MO2-test-R | AATCTTAATAGTGAGCGTGC |
| *Primers for qRT-PCR* | |
| mef2cb-F | ACTCGGACATAGTGGAGACCCTG |
| mef2cb-R | TTCTTTGCAGGCCGTGGTGGG |
| actb1-F | AGATCTTCACTCCCCTTGTTCAC |
| actb1-R | AAACCGGCTTTGCACATACC |
| rpl13a-F | TCTGGAGGACTGTAAGAGGTATGC |
| rpl13a-R | AGACGCACAATCTTGAGAGCAG |

**Table S4. Variants identified in known human CHD genes.**

| Gene | Variant | Mutation | Family | MaF [a] | C score [b] | ClinVar interpretation (N) [ID] |
|---|---|---|---|---|---|---|
| *ANKRD1* | 10-92675322-G-A | p.Ala276Val | 346 | 0.002630 | 23.00 | Benign(5);Likely benign(5);Uncertain significance(1) [45639] |
| *CCDC11* | 18-47765069-C-T | p.Arg407Gln | 3315 | 0.000025 | 25.70 | N/A |
| *CDK13* | 7-40132656-G-A | p.Val1170Met | 3315 | 0.005284 | 26.10 | N/A |
| *CYR61* | 1-86048526-C-G | p.Ser316Cys | 3540 | 0.004719 | 24.60 | N/A |
| *DNAH11* | 7-21600738-A-G | p.Gln311Arg | 226 | 0.000077 | 14.36 | N/A |
| | 7-21760479-G-A | c.7287+5 G>A | 346 | 0.000176 | 14.54 | N/A |
| | 7-21789835-G-A | c.8798-5 G>A | 645 | 0.003884 | 14.13 | Benign(1);Likely benign(2);Uncertain significance(1) [163114] |
| | 7-21788220-C-G | p.Arg2845Gly | 720 | 0.001183 | 16.60 | Likely benign(1);Uncertain significance(1) [359658] |
| | 7-21640338-G-T | p.Glu1015Asp | 1722 | 0.003030 | 22.70 | Benign (2) [238905] |
| *DNAH5* | 5-13850914-C-T | p.Arg1654Gln | 489 | 0.000264 | 32.00 | Uncertain significance (2) [238977] |
| | 5-13914751-C-T | p.Val400Met | 732 | 0.000843 | 34.00 | Likely benign (1) [219733] |
| | 5-13786351-C-G | p.Glu2919Asp | 1121 | 0.003134 | 15.84 | Benign(2);Likely benign(1);Uncertain significance(1) [188365] |
| *EVC2* | 4-5642347-G-C | p.Thr455Arg | 545 | 0.003692 | 23.90 | Benign(3);Likely benign(1);Uncertain significance(1) [193762] |
| | 4-5620263-G-A | p.Ala803Val | 1151 | 0.003871 | 16.90 | N/A |
| *FOXH1* | 8-145700381-C-G | p.Ser113Thr | 346 | 0.004014 | 22.70 | Benign/Likely benign (5) [94385] |
| *KMT2A* | 11-118352769-G-A | p.Ser1325Asn | 49 | 0.000935 | 19.46 | N/A |
| *KMT2D* | 12-49426025-G-T | p.Pro4155Thr | 346 | 0.000028 | 14.07 | N/A |
| | 12-49431844-G-A | p.Arg3099Cys | 3505 | 0.000025 | 24.80 | N/A |

| | | | | | | |
|---|---|---|---|---|---|---|
| *MYH11* | 16-15797967-T-A | p.Thr1934Ser | 732 | 0.001369 | 16.85 | Likely benign(5);Uncertain significance(1) [201044] |
| *MYH6* | 14-23866451-G-T | Leu700Ile | 1710 | N/A | 24.20 | N/A |
| *NFATC1* | 18-77246538-G-A | p.Gly795Arg | 645 | 0.003160 | 14.95 | N/A |
| *NOTCH2* | 1-120468201-A-T | p.Leu1413His | 1121 | 0.003277 | 15.52 | Benign/Likely benign (5)[134975] |
| *NPHP4* | 1-6012895-G-A | c.675 C>T (splice) | 489 | 0.000096 | 5.49 | N/A |
| *PACS1* | 11-66008953-G-A | Gly829Ser | 1151 | N/A | 23.10 | N/A |
| *PDGFRA* | 4-55161324-C-T | p.Thr1052Met | 1151 | 0.000175 | 25.20 | Uncertain significance (2) [39618] |
| *PKD2* | 4-88967919-T-G | p.Phe482Cys | 545 | 0.002039 | 24.90 | Benign (3) [219481] |
| *ROR2* | 9-94487187-C-T | p.Arg530Gln | 732 | 0.001928 | 20.90 | Likely benign (4) [284633] |
| *TBX5* | 12-114837364-T-C | p.Ile106Val | 226 | 0.000999 | 21.80 | Likely benign (3) [213825] |
| *VCAN* | 5-82868257-A-G | p.Asn3253Ser | 346 | 0.000207 | 26.10 | N/A |
| | 5-82816284-T-C | p.Val720Ala | 732 | 0.002033 | 0.001 | Likely benign (3) [354403] |

[a] Minor allele frequency in Genome Aggregation population (123,136 exomes, gnomad.broadinstitute.org). [b] C score according to the Combined Annotation-Dependent Depletion prediction algorithm. N/A: not available.

**Table S5.** Significant protein-protein interaction clusters with at least two CDGs. Statistical significance was determined by permutation analysis (k=10,000). *:P<0.05, **:P<0.01.

| Cluster id. | Cluster size | Mut.Genes | p-value |
|---|---|---|---|
| cluster15 | 27 | 11 | 0.00331 ** |
| cluster17 | 5 | 3 | 0.03872 * |
| cluster215 | 21 | 7 | 0.05427 |
| cluster28 | 13 | 5 | 0.05775 |
| cluster40 | 6 | 3 | 0.06705 |
| cluster202 | 6 | 3 | 0.06807 |
| cluster200 | 12 | 4 | 0.13646 |
| cluster14 | 23 | 6 | 0.19057 |
| cluster8 | 5 | 2 | 0.20428 |
| cluster3 | 29 | 7 | 0.22072 |
| cluster208 | 10 | 3 | 0.24048 |
| cluster10 | 6 | 2 | 0.27304 |
| cluster6 | 6 | 2 | 0.27311 |
| cluster47 | 6 | 2 | 0.27341 |
| cluster230 | 6 | 2 | 0.27429 |
| cluster42 | 6 | 2 | 0.27477 |
| cluster41 | 6 | 2 | 0.27558 |
| cluster13 | 7 | 2 | 0.34492 |
| cluster7 | 8 | 2 | 0.41017 |
| cluster5 | 15 | 3 | 0.48476 |
| cluster4 | 27 | 5 | 0.5071 |
| cluster9 | 17 | 3 | 0.57995 |
| cluster2 | 18 | 3 | 0.61678 |
| cluster113 | 14 | 2 | 0.71902 |
| cluster1 | 30 | 4 | 0.78055 |

**Supplemental Table 6. Rare calcium signaling gene variants shared among affected individuals in multiplex CHD families.**

| Family | Gene | Gene pLI [a] | Variant | Mutation | Confirmed (Sanger seq) | MaF [b] | CADD score [c] | MPC score | Heart malformations |
|---|---|---|---|---|---|---|---|---|---|
| 333 | *ITPR1*[e] | 1.000 | 3-4704816-G-A | V479I | YES | 0.004674 | 22.2 | 1.77 | PAPVR, ASDsv, VSD, COA |
| 346 | *ADCY2*[f] | 1.000 | 5-7695981-G-A | IVS6+5 G>A | YES | 0.004870 | 15.1 | n/a | VSD |
| | *NFAT5*[g] | 1.000 | 16-69660329-A-G | K51E | YES | 0.000031 | 27.0 | 1.95 | VSD |
| 489 | *PLCB2*[h] | 0.000 | 15-40584565-C-A | K802N | YES | 0.000004 | 29.7 | n/a | ASD, subAS, VSD, PDA, HCM |
| | *PLCB2*[h] | 0.000 | 15-40584570-T-A | M801L | YES | n/a | 24.9 | n/a | ASD, subAS, VSD, PDA, HCM |
| | *PLCB2*[h] | 0.000 | 15-40584571-C-A | E800D | YES | n/a | 21.1 | n/a | ASD, subAS, VSD, PDA, HCM |
| 645 | *CACNA1D*[i] | 1.000 | 3-53707750-C-T | A376V | YES | 0.000012 | 32.0 | 2.04 | ASD, VSD |
| 702 | *CACNA1S*[i] | 0.000 | 1-201009430-G-T | P1767T | YES | 0.000244 | 3.34 | 0.11 | VSD, AVSD |
| 732 | *CACNA1H*[i] | 0.756 | 16-1260783-G-A | IVS20-4 G>A | YES | 0.003505 | 22.9 | n/a | TOF |
| | *CACNA1H*[i] | 0.756 | 16-1270119-C-T | R2063W | YES | 0.000078 | 0.07 | n/a | TOF |
| | *GRIA4*[j] | 0.989 | 11-105776016-G-C | G383R | YES | 0.000004 | 26.5 | n/a | TOF |
| 1121 | *CACNA1F*[i] | 0.874 | X-49088240-T-C | T59A | YES | 0.000045 | 0.12 | 0.54 | ASD, VSD, PVS |
| 1151 | *CACNA1S*[i] | 0.000 | 1-201009011-C-T | S1857N | YES | 0.002083 | 23.3 | 0.17 | ASD, EbA |
| 1722 | *ADCY5*[f] | 0.990 | 3-123046510-C-G | E634D | YES | 0.004297 | 15.1 | n/a | ASD, COA, DCM |
| | *ITPR1*[e] | 1.000 | 3-4842276-G-A | A2352T | YES | 0.000809 | 23.5 | 0.92 | ASD, COA, DCM |
| 2077 | *CACNA1I*[i] | 1.000 | 22-40059828-G-C | Q1193H | YES | 0.003845 | 23.7 | n/a | PVS, InfPS |

[a] Probability of loss of function intolerance of gene. [b] Minor allele frequency in gnomAD population (123,136 individuals, http://gnomad.broadinstitute.org). n/a: not available in database.[c] C score according to the Combined Annotation-Dependent Depletion (CADD) prediction algorithm. [d] MPC-score: score based on regional missense constraint. [e] IP3 receptor. [f] Adenylate cyclase. [g] Transcription factor. [h] Phospholipase C. [i] Voltage Dependent Calcium Channel. [j] Glutamate receptor.

**Table S7.** Gene ontology term enrichment of 27 genes within the cluster shown in Figure 3A. P-values were adjusted for multiple testing.

| **Gene Ontology – Molecular Function** | **p-value** |
|---|---|
| calcium-dependent protein kinase C activity (GO:0004698) | 1.61E-06 |
| calcium-dependent protein serine/threonine kinase activity (GO:0009931) | 3.35E-05 |
| calcium-dependent protein kinase activity (GO:0010857) | 4.35E-05 |
| protein kinase C activity (GO:0004697) | 1.47E-04 |
| histone kinase activity (H3-T6 specific) (GO:0035403) | 9.01E-04 |
| protein serine/threonine kinase activity (GO:0004674) | 2.60E-03 |
| ATP binding (GO:0005524) | 2.99E-03 |
| adenyl ribonucleotide binding (GO:0032559) | 3.44E-03 |
| adenyl nucleotide binding (GO:0030554) | 3.57E-03 |
| purine ribonucleoside triphosphate binding (GO:0035639) | 1.03E-02 |
| purine ribonucleoside binding (GO:0032550) | 1.06E-02 |
| purine nucleoside binding (GO:0001883) | 1.07E-02 |
| ribonucleoside binding (GO:0032549) | 1.08E-02 |
| cAMP-dependent protein kinase activity (GO:0004691) | 1.10E-02 |
| histone threonine kinase activity (GO:0035184) | 1.10E-02 |
| nucleoside binding (GO:0001882) | 1.10E-02 |
| purine ribonucleotide binding (GO:0032555) | 1.18E-02 |
| purine nucleotide binding (GO:0017076) | 1.23E-02 |
| ribonucleotide binding (GO:0032553) | 1.31E-02 |
| cyclic nucleotide-dependent protein kinase activity (GO:0004690) | 1.82E-02 |
| protein kinase activity (GO:0004672) | 2.18E-02 |
| adenylate cyclase activity (GO:0004016) | 2.25E-02 |
| carbohydrate derivative binding (GO:0097367) | 3.51E-02 |
| **Gene Ontology – Biological Process** | **p-value** |
| activation of phospholipase C activity (GO:0007202) | 1.59E-07 |
| positive regulation of phospholipase C activity (GO:0010863) | 3.30E-07 |
| regulation of phospholipase C activity (GO:1900274) | 3.83E-07 |
| positive regulation of phospholipase activity (GO:0010518) | 1.03E-06 |
| regulation of phospholipase activity (GO:0010517) | 1.82E-06 |
| positive regulation of lipase activity (GO:0060193) | 2.14E-06 |
| regulation of lipase activity (GO:0060191) | 4.61E-06 |
| activation of protein kinase A activity (GO:0034199) | 3.76E-04 |
| epidermal growth factor receptor signaling pathway (GO:0007173) | 8.48E-04 |
| fibroblast growth factor receptor signaling pathway (GO:0008543) | 8.75E-04 |
| regulation of body fluid levels (GO:0050878) | 1.01E-03 |
| ERBB signaling pathway (GO:0038127) | 1.02E-03 |
| cellular response to fibroblast growth factor stimulus (GO:0044344) | 1.27E-03 |
| response to fibroblast growth factor (GO:0071774) | 1.38E-03 |
| peptidyl-serine phosphorylation (GO:0018105) | 2.19E-03 |

| | |
|---|---|
| neurotrophin TRK receptor signaling pathway (GO:0048011) | 2.37E-03 |
| neurotrophin signaling pathway (GO:0038179) | 2.46E-03 |
| transmembrane receptor protein tyrosine kinase signaling pathway (GO:0007169) | 2.51E-03 |
| histone H3-T6 phosphorylation (GO:0035408) | 2.74E-03 |
| renal water homeostasis (GO:0003091) | 2.74E-03 |
| peptidyl-serine modification (GO:0018209) | 3.18E-03 |
| energy reserve metabolic process (GO:0006112) | 3.27E-03 |
| cellular response to glucagon stimulus (GO:0071377) | 3.86E-03 |
| activation of adenylate cyclase activity (GO:0007190) | 4.52E-03 |
| positive regulation of adenylate cyclase activity (GO:0045762) | 8.41E-03 |
| response to glucagon (GO:0033762) | 8.41E-03 |
| enzyme linked receptor protein signaling pathway (GO:0007167) | 9.95E-03 |
| synaptic signaling (GO:0099536) | 1.11E-02 |
| synaptic transmission (GO:0007268) | 1.11E-02 |
| trans-synaptic signaling (GO:0099537) | 1.11E-02 |
| multicellular organismal water homeostasis (GO:0050891) | 1.48E-02 |
| positive regulation of cyclase activity (GO:0031281) | 1.72E-02 |
| positive regulation of lyase activity (GO:0051349) | 1.72E-02 |
| water transport (GO:0006833) | 1.90E-02 |
| water homeostasis (GO:0030104) | 1.99E-02 |
| fluid transport (GO:0042044) | 2.49E-02 |
| regulation of adenylate cyclase activity (GO:0045761) | 2.60E-02 |
| histone-threonine phosphorylation (GO:0035405) | 3.35E-02 |
| positive regulation of cAMP biosynthetic process (GO:0030819) | 3.87E-02 |
| immune system process (GO:0002376) | 3.91E-02 |
| cellular response to forskolin (GO:1904322) | 4.38E-02 |
| response to forskolin (GO:1904321) | 4.38E-02 |
| regulation of cyclase activity (GO:0031279) | 4.63E-02 |
| regulation of lyase activity (GO:0051339) | 4.97E-02 |

**Table S8.** Replication using WES data from 714 CHD cases and 4922 controls. The number of cases and controls with rare variants (MAF < 0.001) in any of the genes *ADCY2*, *ADCY5*, *CACNA1D*, *CACNA1H*, *CACNA1I*, *CACNA1S*, *GRIA4*, *ITPR1*, *NFAT5* and *PLCB2* was compared (see supplemental material for details). WES data was not available for *CACNA1F*.

| Type of rare variant [a] | Cases | Controls | P-value [b] | | Odds ratio |
|---|---|---|---|---|---|
| Protein sequence altering variants | 116 | 877 | 3.18E-01 | n.s. | 0.89 [0.72-1.11] |
| Protein altering variants, MPC score >1 | 43 | 243 | 3.35E-01 | n.s. | 1.23 [0.86-1.73] |
| Protein altering variants, MPC score >2 | 21 | 55 | 3.69E-04 | ** | 2.68 [1.53-4.54] |
| Protein truncating variants | 3 | 16 | 7.25E-01 | n.s. | 1.29 [0.24-4.53] |
| Synonymous variants | 94 | 630 | 7.65E-01 | n.s. | 1.03 [0.81-1.31] |

[a] Variants with pathogenic effect was defined by MPC score >2. [b] Fisher's exact test. **:P<0.01. n.s.: not significant.
